# Mathematical modelling of a novel bioactive glass treatment for bacterial biofilms

**DOI:** 10.64898/2026.08.10.743863

**Authors:** Sandeep Shirgill, Sarah A. Kuehne, Gowsihan Poologasundarampillai, Sara Jabbari, John Ward

## Abstract

Chronic wounds (principally pressure sores, venous leg ulcers and diabetic foot ulcers) are a drain on global health services and remain a major area of unmet clinical need. Chronic wounds are characterised by a bacterial biofilm (densely aggregated colonies of bacteria encased by a matrix of extracellular polymeric substances), which hinders innate immune response and can prevent wound healing. Bioactive glass (BG) fibres doped with antimicrobial metal ions, such as silver, can offer a promising treatment for chronic wound infections, where silver is well known for its antimicrobial activity against a range of pathogens and is commonly used in wound dressings.

We first present a system of non-linear partial differential equations to model the treatment of a chronic wound biofilm infection with BG fibres. The BG fibres are assumed to have two mechanisms of action against the biofilm: physical disruption of the top layers of the biofilm by the BG fibres; and release of antimicrobial silver ions from the BG fibres, which then diffuse into the biofilm and can kill the bacteria. Treatment-associated parameters are estimated from *in vitro* experimental data using a combination of least-squares minimisation and Approximate Bayesian Computation (ABC). Sensitivity investigations are performed on other parameters that cannot currently be calculated experimentally to investigate their influence on treatment efficacy. We thus predict key parameter regimes that should lead to biofilm eradication, crucially informing the future design of metal-doped BG fibres to maximise treatment efficacy.

**Author summary:** Chronic wounds are a huge drain on global health services and will become even more problematic due to an ageing population. Current treatment methods are often unsuccessful, where treatment failure is exacerbated by the presence of a biofilm infection. Biofilms consist of communities of bacteria that adhere to the wound surface and produce extracellular polymeric substances, which can protect the bacteria by acting as both a physical and chemical barrier. More recently, there has been a focus on biofilm-based wound care, where the aim is to firstly eradicate the biofilm infection, which then enables wound healing to occur naturally. Our aim is to produce a novel treatment that can target and eradicate the biofilm infection, followed by directly assisting the wound healing. Bioactive glass (BG) fibres doped with silver offer a promising treatment as they have both anti-biofilm effects and can also stimulate the wound healing process. Here, we restrict attention to their anti-biofilm properties. By developing a mathematical model, we can predict treatment outcomes under several different scenarios, the results of which can then be utilised during design of the BG fibres. Using this combination of computational and experimental approaches, we reduce both the cost and time of optimising this promising treatment.

## Introduction

Chronic wound infections represent a pressing challenge for global health systems, with conditions like pressure sores, venous leg ulcers, and diabetic foot ulcers imposing substantial burdens. These wounds not only incur significant healthcare costs, estimated at £8.3 billion annually in the United Kingdom’s National Health Service alone (based on data from 2017/2018), but also present a major unmet clinical need [1, 2]. As the prevalence of diabetes and the ageing population continue to increase, the burden and incidence of chronic wounds are expected to rise, highlighting the urgent need for more effective treatment strategies.

One potential avenue for addressing chronic wound infections lies in biofilm-based wound care. Biofilms, comprising densely aggregated bacterial colonies encased in extracellular polymeric substances (EPS), are prevalent in chronic wounds, contributing to delayed healing and treatment resistance [3–6]. In addition to experimental evidence, mathematical modelling has shown that biofilms substantially impair wound healing and that effective biofilm-targeted interventions can accelerate healing [7], supporting strategies that directly disrupt or eradicate biofilms [5, 8]. However, biofilms exhibit heightened antimicrobial tolerance compared to planktonic bacteria (free-floating cells), posing a significant challenge to treatment efficacy [9]. Fig 1 illustrates the main mechanisms for reduced susceptibility against antimicrobial treatment, namely:

1. Impairment of antimicrobial diffusion throughout the biofilm;
2. Physiological heterogeneity in the bacterial population due to substrate limitation in the biofilm;
3. The presence of sub-populations of persister cells in the biofilm that evade antimicrobial effects [10];
4. Bacteria employing an adaptive response, e.g. the use of efflux pumps or the production of antibiotic-degrading enzymes [9].

**Fig 1.**
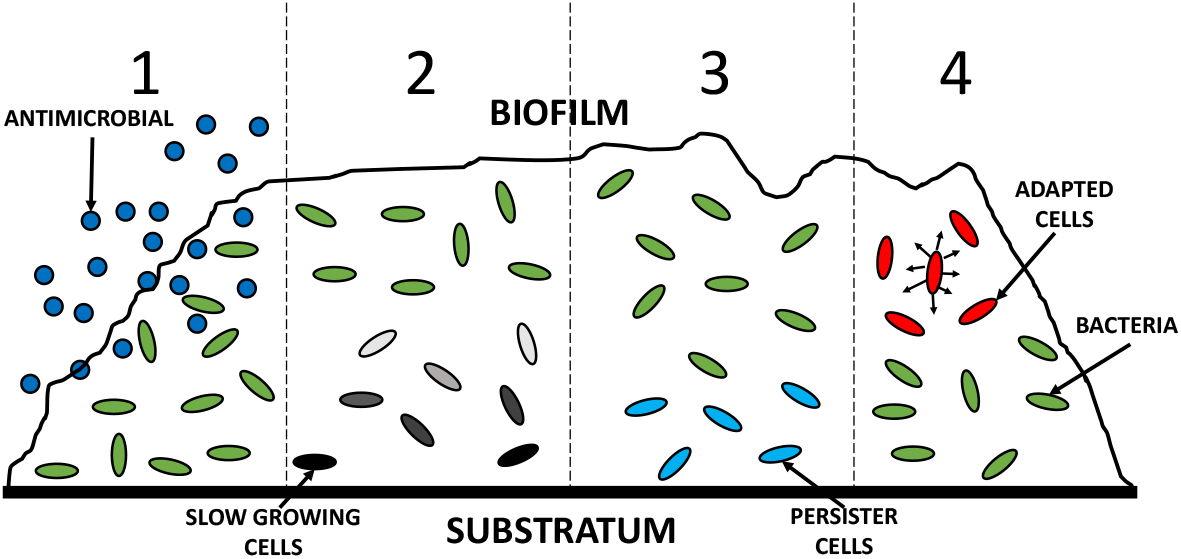
Four main mechanisms of reduced susceptibility of biofilms to antimicrobial adapted from [9]. 1 - antimicrobial concentration gradients through difficulty of diffusion. 2 - gradients of metabolic substrates, creating regions of slow growing bacteria near the bottom of the biofilm that are more tolerant to antimicrobial effects. 3 - the formation of dormant persister cells that evade antimicrobial action and revert back to normal cells and resume growth. 4 - employment of an adaptive response.

In this context, bioactive glass (BG) fibres emerge as a promising treatment modality. These fibres possess both wound healing and antimicrobial properties, with silver-doped variants demonstrating particularly potent activity against antibiotic-resistant strains and biofilms [11–17]. Utilising the antimicrobial capabilities of BG fibres holds potential for combating biofilm infections in chronic wounds.

To gain deeper insights into the dynamics of biofilm growth and treatment generally, mathematical models have been explored. These models elucidate the complex interactions within biofilms and their response to antimicrobial interventions [9, 18–26]. Ranging from discrete to continuum approaches, these models provide valuable tools for understanding biofilm behaviour and optimising treatment strategies.

Our study contributes to this body of research by developing a mathematical model to predict the treatment efficacy of BG fibres against biofilm infections in chronic wounds. To facilitate parameterisation using experimental data, we begin by formulating a simplified core model, which provides a foundation for later extensions that incorporate more complex mechanisms. In the Results section, we first describe the parameterisation of substrate-related dynamics (focusing on oxygen and antimicrobial diffusion), before estimating treatment-associated parameters using experimental data. We then use these estimates to explore how key mechanisms (such as physical disruption, ion release, diffusion, and antimicrobial removal) shape treatment outcomes.

The parameter estimation carried out in this study provides critical insights that enhance our understanding of treatment efficacy, guide experimental practice, and inform the development of optimal BG fibre formulations for improved antimicrobial performance. This work advances our understanding of biofilm-based wound care and paves the way for more effective chronic wound treatments. To our knowledge, this study is the first to model the treatment of biofilms with ion-doped fibres, offering a novel framework for future research.

## Methods & Materials

### Experimental materials

The *Pseudomonas aeruginosa* strain, PAO1-N, was kindly provided by Paul Williams, University of Nottingham. Whatman Nuclepore^®^, polycarbonate membranes (0.1 *µ*m pore size, 19 mm diameter) were purchased from Merck^®^. Roswell Park Memorial Institute (RPMI) 1640 was purchased from Gibco™. Silver nitrate (99%+ ACS reagent) was purchased from Acros Organics^®^. Ethanol was purchased from Fisher Scientific™. All other reagents were purchased from Sigma-Aldrich^®^.

### Experimental methods

Bioactive glass fibres, either undoped or doped with 2 mol% silver(I) oxide, were electrospun as described in [17].

To measure the total concentration of silver species released from the fibres during soaking, including both silver ions and silver nanoparticles, silver-doped fibres were placed in 5 mM (pH=7.4) tris buffer in the ratio of 1.5 mg/ml and incubated at 37°C. At specific time points (t=0 mins, 10 mins, 30 mins, 1 hr, 2 hrs, 8 hrs, 1 day, 2 days, 3 days & 7 days), 100 *µ*l aliquots of dissolution products were removed from the sample and diluted 1:100 in RO water. The concentration of silver in the dissolution products was measured by research technician, Christopher Stark, in the Geography, Earth and Environmental Sciences department at the University of Birmingham using the PerkinElmer Nexion 300X model for induced coupled plasma mass spectrometry (ICP-ms).

To investigate the antimicrobial effects of BG fibres on PAO1-N biofilms, nuclepore^®^ polycarbonate membranes were inoculated with an overnight culture of PAO1-N, which was diluted to OD_600_=0.05 and placed in the wells of a 12-well plate containing 1.5 ml of BHI agar. Wells containing inoculated membranes were filled with 1 ml of RPMI 1640 and then incubated for 24 hrs at 37°C, to allow biofilm formation to occur on the membrane. After incubation, the RPMI 1640 was replaced with fresh media and 10 mg of either undoped fibres or silver-doped fibres were added to the wells. After *t* =0, 1, 3, 5, 7 & 24 hrs of treatment, membranes containing the biofilm were then removed from wells, placed in 2 ml of BHI broth, and vortexed for 1 minute to resuspend any biofilm formation. Biofilm viability was assessed using the Miles and Misra method to quantify the CFU/ml [27].

### Modelling assumptions

To isolate antimicrobial transport effects, we model the biofilm as a layer of fixed height [28, 29]. The biofilm is treated as an incompressible, volume-filling mixture with constant water and EPS fractions, following previous continuum formulations [30–32]. Water is included explicitly, as it can constitute approximately 70% of the biofilm mass [33]. Consequently, only the live and dead biomass fractions are permitted to vary within the biofilm, while its total volume is held constant. Extensions to growing biofilms with time-dependent thickness are left for future work. A schematic of the biofilm model is shown in Fig 2 with variable definitions and units provided in Table 1.

**Table 1.** Variable definitions for Eqs (1)-(6).

| Variables | Description | Units |
| --- | --- | --- |
| $\epsilon_x(z, t)$ | Live cell fraction | - |
| $\epsilon_z(z, t)$ | Dead cell fraction | - |
| $O(z, t)$ | Proportion of available oxygen | - |
| $A(z, t)$ | Antimicrobial ion concentration within biofilm | $\mu\text{g ml}^{-1}$ |
| $A_b(t)$ | Antimicrobial ion concentration within bulk fluid | $\mu\text{g ml}^{-1}$ |
| $\phi(z)$ | BG fibre fraction | - |

**Fig 2.**
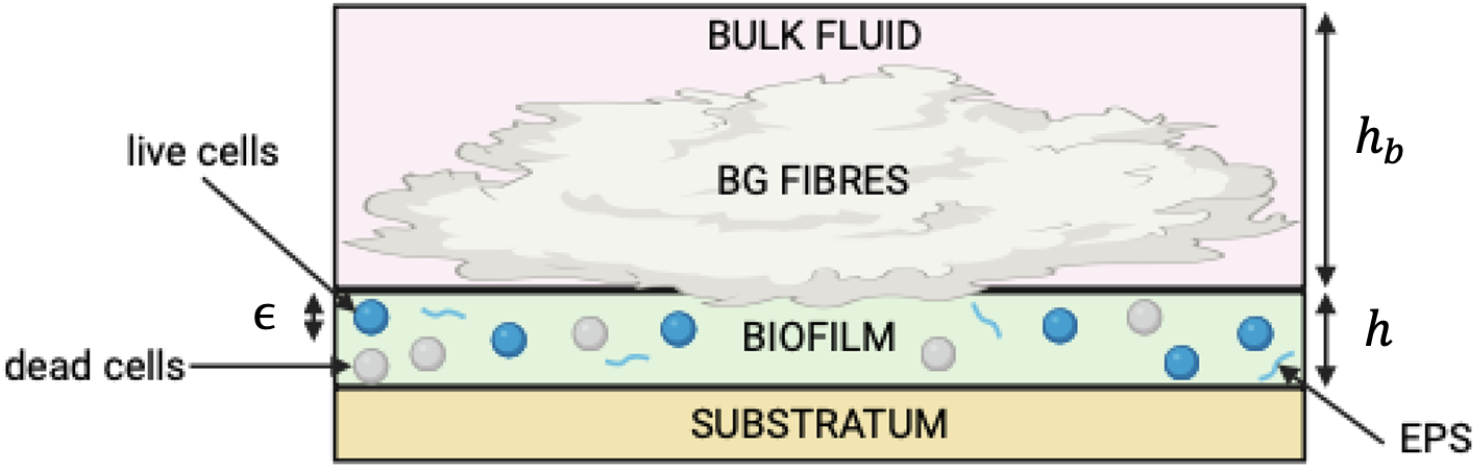
A schematic of the biofilm model. The biofilm forms on top of the substratum and is composed of live cells, dead cells, EPS and water. Fibres are placed in bulk fluid (of height *h*_*b*_) above the biofilm, with a height *h*, and penetrate into the biofilm with length *ϵ*. Created in BioRender.com.

The following further assumptions are then made to formulate the model:

- The biofilm volume consists only of live cells (*ϵ*_*x*_(*z, t*)), dead cells (*ϵ*_*z*_(*z, t*)), EPS (*ϵ*_*b*_), and water (*ϵ*_*w*_), assuming the presence of only one bacterial species. Nutrients are abundant, with oxygen (*O*(*z, t*)) being the sole growth-limiting substrate.
- The biofilm is modelled in one dimension, with variables changing along its depth. It has a height *h* and is attached to an impermeable substratum at *z* = 0. The media above the biofilm has a significantly greater height, denoted as *h*_*b*_ (i.e. *h* ≪ *h*_*b*_).
- Before treatment, the biofilm is considered to be in dynamic equilibrium, where bacterial growth is balanced by natural cell death. Consequently, these processes are not explicitly modelled.
- During treatment, BG fibres (*ϕ*(*z*)) are soaked in the bulk fluid but can also penetrate and disrupt the top layers of the biofilm, with a penetration depth *ϵ*. Antimicrobial (silver) ions are released from the fibres at a constant rate until *t* = *t*_*a*_, with *A*(*z, t*) and *A*_*b*_(*t*) denoting the antimicrobial concentrations within and above the biofilm, respectively. This cut-off approximates experimental release profiles, where ion release is initially approximately linear before plateauing, indicating negligible subsequent release. Removal of antimicrobial ions occurs through interactions with live cells, binding by EPS, and chemical neutralisation by media components (modelled as a sink term).
- Reduced susceptibility of biofilms to antimicrobial treatment is captured through two mechanisms. Firstly, antimicrobial concentration gradients arise due to diffusion limitations (as illustrated in the first mechanism of resistance in Fig 1). Secondly, bacteria deeper within the biofilm experience lower oxygen levels, leading to slower growth and reduced antimicrobial uptake (see the second mechanism of resistance in Fig 1). This physiological resistance mechanism is modelled using Monod kinetics, drawing from concepts outlined by Cogan *et al* (2005) [26].

### Model formulation

From the above assumptions, the following system of partial differential equations (PDEs) are formulated to describe BG fibre treatment of an *in vitro* biofilm model:

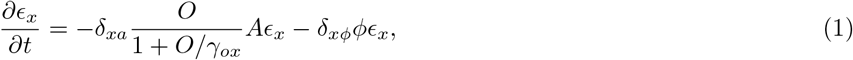

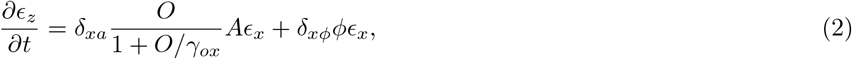

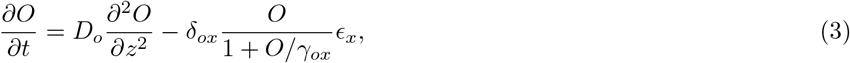

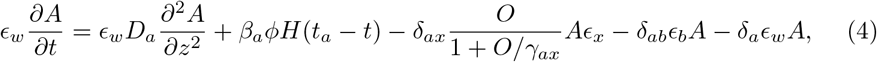

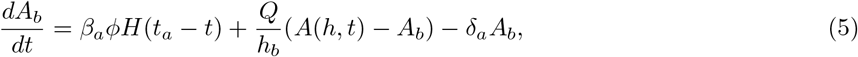

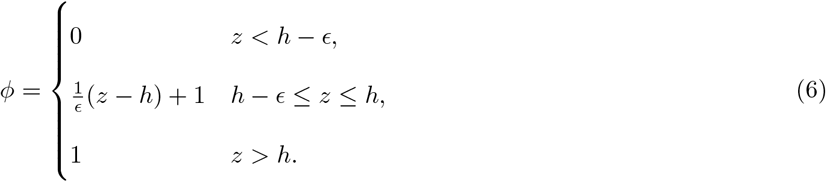

The function *H*(*t*_*a*_ − *t*) in (4) and (5) is the Heaviside step function, defined here as *H*(*t*_*a*_ − *t*) = 1 for *t* ≤ *t*_*a*_ and *H*(*t*_*a*_ − *t*) = 0 for *t* > *t*_*a*_, representing the end-time of ion release. Parameter definitions can be found in Table 2 and detailed derivation for Eqs. (1)-(6) can be found in [34]. From conservation of volume, we have that

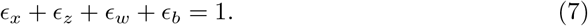

**Table 2.** Parameter definitions for (1)-(6).

| Parameters | Description | Units |
| --- | --- | --- |
| $\epsilon_{x0}$ | initial live cell fraction | - |
| $\epsilon_{z0}$ | initial dead cell fraction | - |
| $\epsilon_w$ | biofilm water fraction | - |
| $\delta_{xa}$ | death rate of live cells by antimicrobial | $\text{ml } \mu\text{g}^{-1} \text{ hr}^{-1}$ |
| $\delta_{x\phi}$ | death rate of live cells by physical disruption of the BG fibres | $\text{hr}^{-1}$ |
| $\gamma_{ax}$ | half-saturation constant for oxygen related to antimicrobial effectiveness | - |
| $\gamma_{ox}$ | half-saturation constant for oxygen related to its consumption by live cells. | - |
| $D_o$ | diffusion coefficient of oxygen | $\mu\text{m}^2 \text{ hr}^{-1}$ |
| $\delta_{ox}$ | uptake rate of oxygen by live cells | $\text{hr}^{-1}$ |
| $D_a$ | diffusion coefficient of antimicrobial ions | $\mu\text{m}^2 \text{ hr}^{-1}$ |
| $\beta_a$ | release rate of antimicrobial ions | $\mu\text{g ml}^{-1} \text{ hr}^{-1}$ |
| $t_a$ | duration of antimicrobial ion release | $\text{hr}$ |
| $\delta_{ax}$ | uptake rate of antimicrobial ions by live cells | $\text{hr}^{-1}$ |
| $Q$ | exchange rate of antimicrobial ions at the biofilm/bulk fluid interface | $\mu\text{m hr}^{-1}$ |
| $h_b$ | height of the bulk fluid | $\mu\text{m}$ |
| $h$ | height of the biofilm | $\mu\text{m}$ |
| $\epsilon$ | depth of penetration by the BG fibres into the biofilm | $\mu\text{m}$ |
| $\delta_a$ | removal rate of ions in the media | $\text{hr}^{-1}$ |
| $\delta_{ab}$ | binding rate of antimicrobial ions to EPS | $\text{hr}^{-1}$ |
| $\epsilon_b$ | biofilm EPS fraction | - |

The boundary conditions are given by,

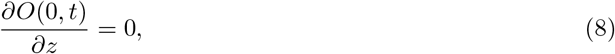

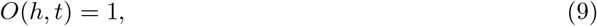

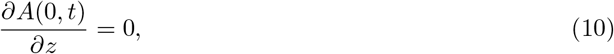

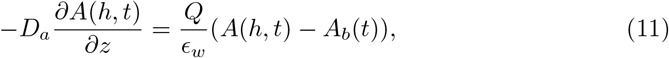

where (8) and (9) represent no oxygen flux at the biofilm/substratum interface and constant oxygen concentration at the top of the biofilm respectively. No flux of antimicrobial ions at the biofilm/substratum interface is represented by (10). Finally (11) captures the flux of ions between the biofilm/bulk liquid interface. The initial conditions for the model are:

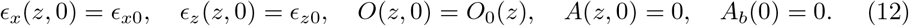

Here, it is assumed that the live and dead cell fractions are constant throughout the biofilm at the start of treatment.

### Parameter estimation techniques

To estimate parameters, we minimise the square of the error (least squares) using the Nelder-Mead simplex algorithm to determine parameter estimates [47]. Secondly, we employ Approximate Bayesian Computation (ABC) to identify the range of possible parameter estimates that produce suitable predictions [48].

The least squares method offers a point estimate, whereas the ABC method provides a range of parameter estimates, yielding a more comprehensive understanding of model performance and fit to experimental data. By utilising both approaches, we not only obtain a point estimate but also assess the robustness and uncertainty associated with the parameter estimates, thereby enhancing the reliability and interpretability of the model. This dual approach ensures a more thorough exploration of parameter space and improves overall confidence in the model’s predictions.

## Results and Discussion

We estimate parameters in Eqs. (1)-(6) through empirical observations wherever feasible, resorting to established values from the literature or imposing values that produce qualitatively realistic behaviours when direct experimental data is unavailable. Initially, our analysis focuses solely on antimicrobial elimination resulting from interactions with live cells, presuming negligible binding to EPS and absence of neutralisation, thus setting *δ*_*a*_ = *δ*_*ab*_ = 0. This simplification allows us to develop and parameterise a more tractable model using experimental data before incorporating additional mechanisms of antimicrobial removal. Furthermore, we assume negligible EPS presence, setting *ϵ*_*b*_ = 0. This simplification streamlines the parameterisation process. Subsequent sections delve into the intricate dynamics of EPS binding and neutralisation of the antimicrobial ions.

### Substrate interactions within biofilms

In this section, we focus on parameterising substrate-related dynamics, beginning with oxygen. Oxygen behaviour in biofilms is well-characterised in the literature [28, 35–44], making it a suitable starting point for model development and validation. This also allows us to assess whether simplifications, such as the thin biofilm assumption, are appropriate based on biofilm thickness. Parameters associated with oxygen, including diffusion and utilisation by bacteria, are provided in Table 7.

Our timescale of interest is the period in which cell death is caused by antimicrobial action, which is *T* ∼ 1 hr. Given this (using parameter values from Table 7), *δ*_*ox*_*T* ≫ 1 and 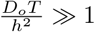 provided *h* ≪ 2000 *µ*m, which is true for typical chronic wound biofilms [45]. Therefore, oxygen can be assumed to be in a quasi-steady state, simplifying the PDE for oxygen, (3), to:

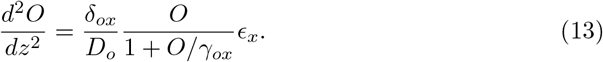

From [43], *γ*_*ox*_ ≈ 0.0417 ≪ *O*(*h, t*) = 1 so for thin biofilms we expect *γ*_*ox*_ ≪ *O*(*z*) for 0 < *z* ≤ *h*, allowing Eq. (13) to be further simplified to:

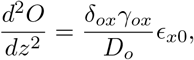

which can be solved to give:

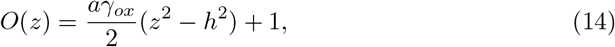

where 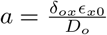.

Based on literature and previous experimental work, it is known that *ϵ*_*x*0_ ≈ 0.3 [33, 34], giving *a* ≈ 0.012. To determine the biofilm height at which the thin biofilm assumption holds, we compare the solution to the quasi-steady ordinary differential equation (ODE) for oxygen, (13), with solutions obtained using the assumption *O* ≫ *γ*_*ox*_, (14), at various heights. Table 3 shows the proportion of oxygen at the bottom of the biofilm for various heights. For biofilms up to 20 *µ*m thick, solutions using the thin biofilm assumption are similar to those from the quasi-steady ODE. Thus, we use the thin biofilm assumption for biofilms ≤ 20 *µ*m thick.

**Table 3.** Solution for *O*_0_(*z*) **at** *z* = 0, for different biofilm heights, *h*, comparing solutions to (13) to solutions assuming *O* ≫ *γ*_*ox*_.

| **h ($\mu\text{m}$)** | **$O \gg \gamma_{ox}$ (14)** | **ODE solution (13)** |
| --- | --- | --- |
| 10 | 0.975 | 0.976 |
| 20 | 0.900 | 0.904 |
| 50 | 0.375 | 0.423 |
| 100 | -1.502 | 0.0018 |

For biofilms 50 *µ*m thick, the thin biofilm assumption starts to diverge from the quasi-steady ODE solutions, indicating it breaks down at this height. For 100 *µ*m thick biofilms, the thin biofilm assumption predicts negative oxygen levels at the base, making the solutions unphysical and invalid at these heights. This confirms that the thin biofilm assumption is not appropriate for biofilms of these thicknesses.

Solutions to (13) with the given parameters align well with the literature. Previous studies suggest oxygen penetration in biofilms is about 77 *µ*m under steady-state conditions with zero-order kinetics, consistent with experimental data showing negligible oxygen at the base of 100 *µ*m biofilms [28, 37]. While our focus is on oxygen distribution in thin biofilms, the parameter values can be extrapolated to thicker biofilms, where *O*(*z*) ≪ *γ*_*ox*_.

Our experimental observations indicate that biofilms grown for this study are typically around 20 *µ*m thick, allowing us to apply the thin biofilm assumption. This simplifies the oxygen dynamics while remaining consistent with the observed characteristics of chronic wound biofilms. As a result, we use the simplified solution for oxygen, given by Eq. (14), throughout the remainder of the study.

We next consider ion diffusion of silver ions within the biofilm. To estimate *D*_*a*_, we start with the assumption that electrostatic attraction between negatively charged EPS and silver has minimal impact on silver transport within the biofilm matrix. Additionally, we consider whether silver is released directly as ions or as nanoparticles that diffuse into the biofilm before releasing ionic silver. Stokes-Einstein law [46] for small spheres tells us that the diffusion rate *D* ∝ 1*/r*, where *r* is the radius of the sphere; using this as a basis, it follows that *r*_*a*_*D*_*a*_ = *r*_*o*_*D*_*o*_, hence *D*_*a*_ = *r*_*o*_*D*_*o*_*/r*_*a*_. In the ionic form of silver this leads to *D*_*a*_ ≈ 6.5 × 10^6^ *µ*m^2^ hr^−1^ and for the (much larger) nanoparticle form *D*_*a*_ ≈ 7.5 × 10^3^ *µ*m^2^ hr^−1^. In our analysis we will use the latter value as the default parameter unless stated otherwise, which helps maintain methodological consistency and reliability across the analysis. Consequently, similar to oxygen, antimicrobial diffusion occurs at a significantly faster rate compared to the timescale of the model, typically spanning hours. We therefore apply a quasi-steady-state assumption for *A* to effectively capture its dynamics,

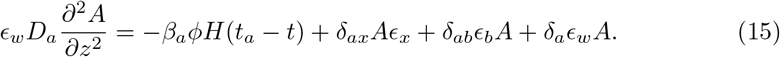

### Parameter estimation: ion release from BG fibres

The parameters *β*_*a*_ (ion release rate) and *t*_*a*_ (duration of ion release) are estimated from experimental data measuring the release of silver species from BG fibres over time when soaked in media (shown in Fig 3). In 8 hours, there was a burst release of silver species from 0 *µ*g ml^−1^ to approximately 6.5 *µ*g ml^−1^. Therefore we set *t*_*a*_ = 8 and estimate *β*_*a*_ ≈ 6.5*/*8 *µ*g ml^−1^ hr^−1^.

**Fig 3.**
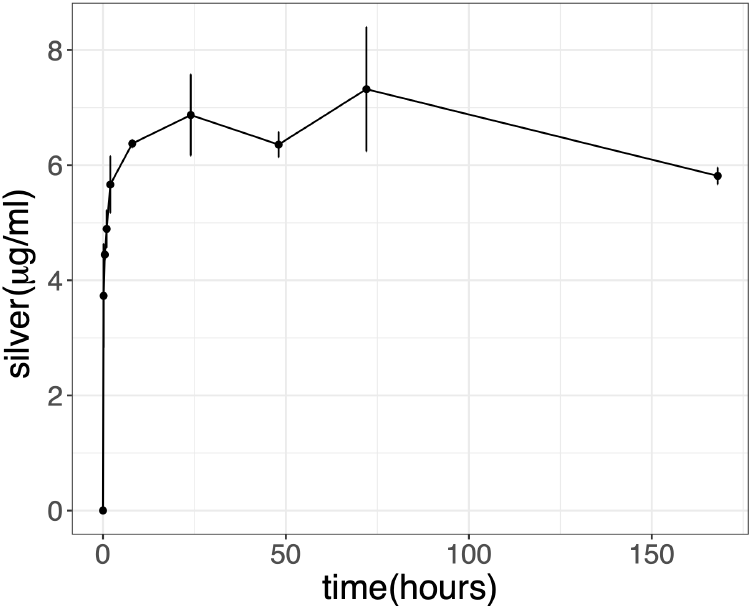
Experimental data: Silver species concentration released from silver-doped BG fibres soaked in media over time. The error bars indicate the standard deviation of the data (n=3).

### Parameter estimation: Interactions between BG fibres and released ions with cells

We use experimental data to measure *δ*_*xa*_ (death rate by antimicrobial), *δ*_*ax*_ (uptake rate of antimicrobial), and *δ*_*xϕ*_ (death rate by physical disruption) (see Fig 4a) using a combination of two approaches.

**Fig 4.**
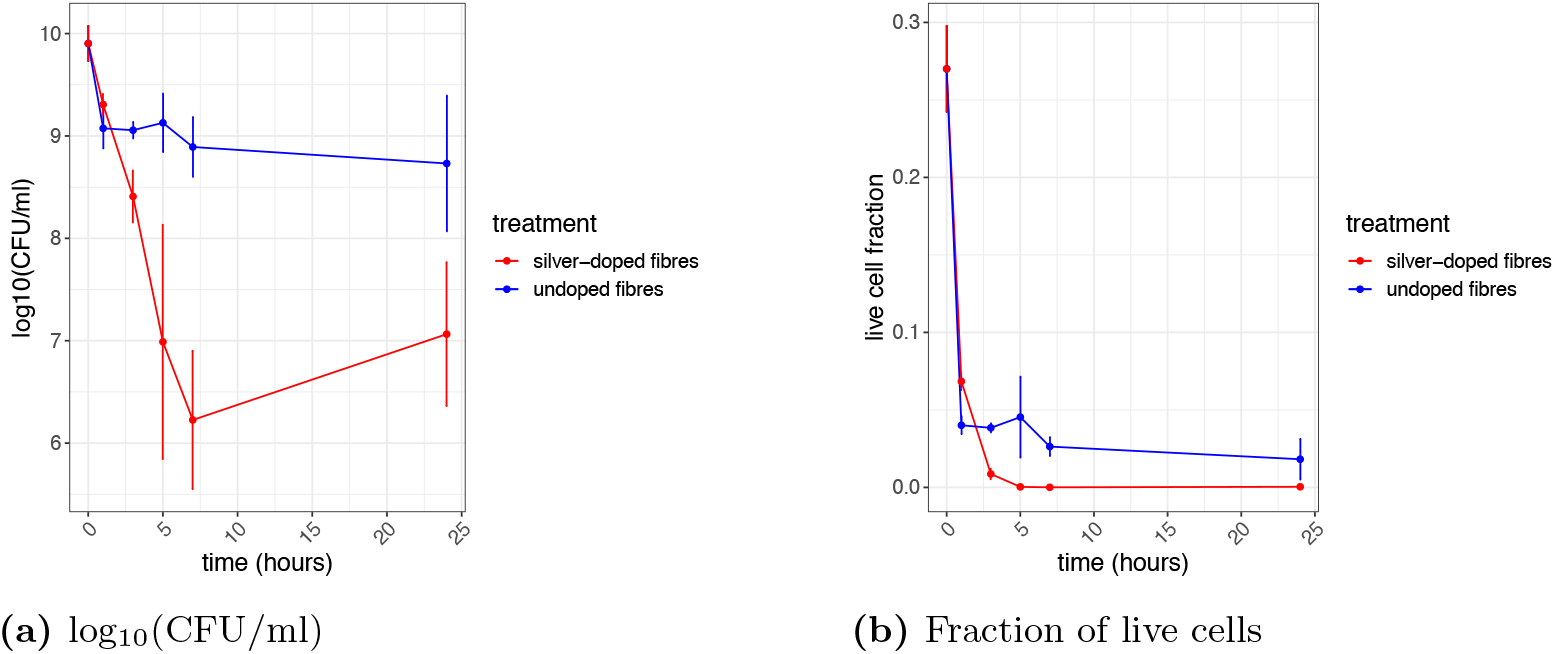
Experimental data: Treatment of *P. aeruginosa* biofilms with undoped fibres or silver-doped fibres measured over 24hrs. Data is converted from log_10_(CFU/ml) (Fig 4a) to fraction of live cells, *ϵ*_*x*_ (4b). The error bars indicate the standard deviation of the data (n=3).

Firstly, assumptions are made to simplify the model (1)-(6) and facilitate parameter estimates in this section:

- From our experimental data, biofilms had an average height of approximately 20 *µ*m. Biofilms at this height have been shown to have a negligible reduction of oxygen within the biofilm (see Table 3), therefore we can assume that *O* ≫ *γ*_*ox*_.
- Furthermore, given the thinness of the biofilms, it is reasonable to assert that there is minimal spatial variation, allowing for complete diffusion of ions within the biofilm. Therefore, the spatial gradient of ions can be considered negligible, effectively reducing the governing PDEs to ODEs.
- Notably, ion exchange at the interface between the biofilm and bulk fluid is disregarded, under the assumption that ions are instantaneously released into the biofilm.

From these assumptions, the reduced biofilm model becomes,

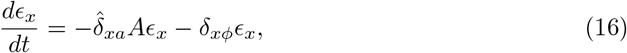

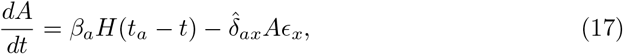

where 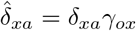 and 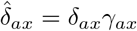. In the rest of this section, the hats are dropped for ease of notation. Note that (as already described) terms describing antimicrobial removal through interactions with EPS and neutralisation are not included as we assume that antimicrobial removal through interactions with live cells is the dominating effect.

#### Determining *δ*_*xϕ*_

To estimate the physical disruption rate inflicted on bacteria by the BG fibres (*δ*_*xϕ*_), we utilise data from the treatment of biofilms by BG fibres that are not doped with the antimicrobial. In the absence of antimicrobial ions the ODEs, (16)-(17), reduce to the following:

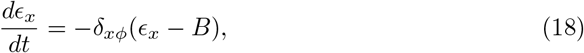

where *B* signifies the proportion of live cells remaining after disruption and may be inferred from the penetration depth of fibres into the biofilm, *ϵ*. This extra parameter is required to prevent *ϵ*_*x*_ from converging to zero, as not all cells die after treatment. It also helps capture physical disruption primarily occurring in the top portion of the biofilm.

Equation (18) can be solved analytically to obtain,

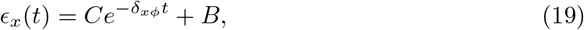

where *C* + *B* is the initial proportion of live cells within the biofilm (i.e. *ϵ*_*x*0_ = *C* + *B*). Here, *C* and *B* are given initial values but are also included as fitted parameters.

To fit the parameters in Eq. (19) to experimental data, the data in Fig. 4b (death rate of biofilms subjected to fibres) are scaled to represent the fraction of live cells remaining, rather than CFU/ml. Using this scaled data, the parameters in Eq. (19) are estimated based on the experimental results from biofilm treatment with undoped fibres to isolate the effects of physical disruption from antimicrobial action (represented by the blue line in Fig 4b with values provided in Table 4).

**Table 4.**
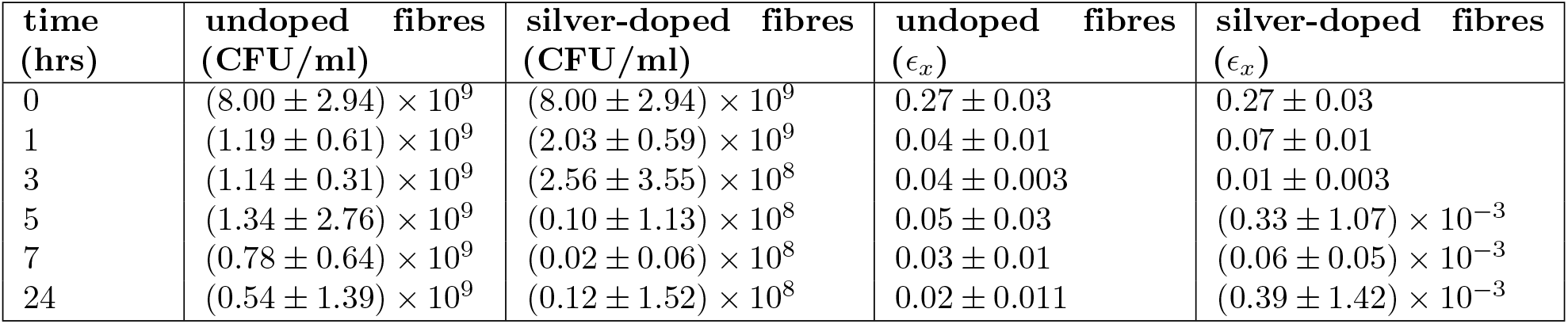
Biofilm viability (measured in CFU/ml) and corresponding calculated values of live cell fractions at timepoints when treated with either undoped or silver-doped fibres.

| time<br>(hrs) | undoped fibres<br>(CFU/ml) | silver-doped fibres<br>(CFU/ml) | undoped fibres<br>($\epsilon_x$) | silver-doped fibres<br>($\epsilon_x$) |
| --- | --- | --- | --- | --- |
| 0 | $(8.00 \pm 2.94) \times 10^9$ | $(8.00 \pm 2.94) \times 10^9$ | $0.27 \pm 0.03$ | $0.27 \pm 0.03$ |
| 1 | $(1.19 \pm 0.61) \times 10^9$ | $(2.03 \pm 0.59) \times 10^9$ | $0.04 \pm 0.01$ | $0.07 \pm 0.01$ |
| 3 | $(1.14 \pm 0.31) \times 10^9$ | $(2.56 \pm 3.55) \times 10^8$ | $0.04 \pm 0.003$ | $0.01 \pm 0.003$ |
| 5 | $(1.34 \pm 2.76) \times 10^9$ | $(0.10 \pm 1.13) \times 10^8$ | $0.05 \pm 0.03$ | $(0.33 \pm 1.07) \times 10^{-3}$ |
| 7 | $(0.78 \pm 0.64) \times 10^9$ | $(0.02 \pm 0.06) \times 10^8$ | $0.03 \pm 0.01$ | $(0.06 \pm 0.05) \times 10^{-3}$ |
| 24 | $(0.54 \pm 1.39) \times 10^9$ | $(0.12 \pm 1.52) \times 10^8$ | $0.02 \pm 0.011$ | $(0.39 \pm 1.42) \times 10^{-3}$ |

First, for the least squares method, we initiate the parameter estimation process by selecting initial guesses for parameter values. These initial guesses allow us to generate the first iteration of model solutions, which can then be compared to the experimental data. Our initial guess for *δ*_*xϕ*_ is 3 hr^−1^. For the initial conditions, we set *ϵ*_*x*0_ = *C* + *B* = 0.27 as our starting guess. This value corresponds to the initial experimental data point at *t* = 0 hrs, where there were 8 × 10^9^ CFU/ml within the biofilm. By establishing this relationship, we convert all other values in the data set to the live cell fraction (as shown in Fig. 4b). To estimate the initial guess for parameter *B*, we consider the value of *ϵ*_*x*_ at *t* = 24 hrs in the data set, which is approximately 0.02.

Taking these initial guesses together, we employ the built-in MATLAB optimizer *fminsearch()* to minimise the least squares error, giving an estimate for a default value of *δ*_*xϕ*_ = 3.81 hr^−1^. Fig 5a illustrates the good agreement between the solution to Eq (19) and the data with this choice of *δ*_*xϕ*_.

**Fig 5.**
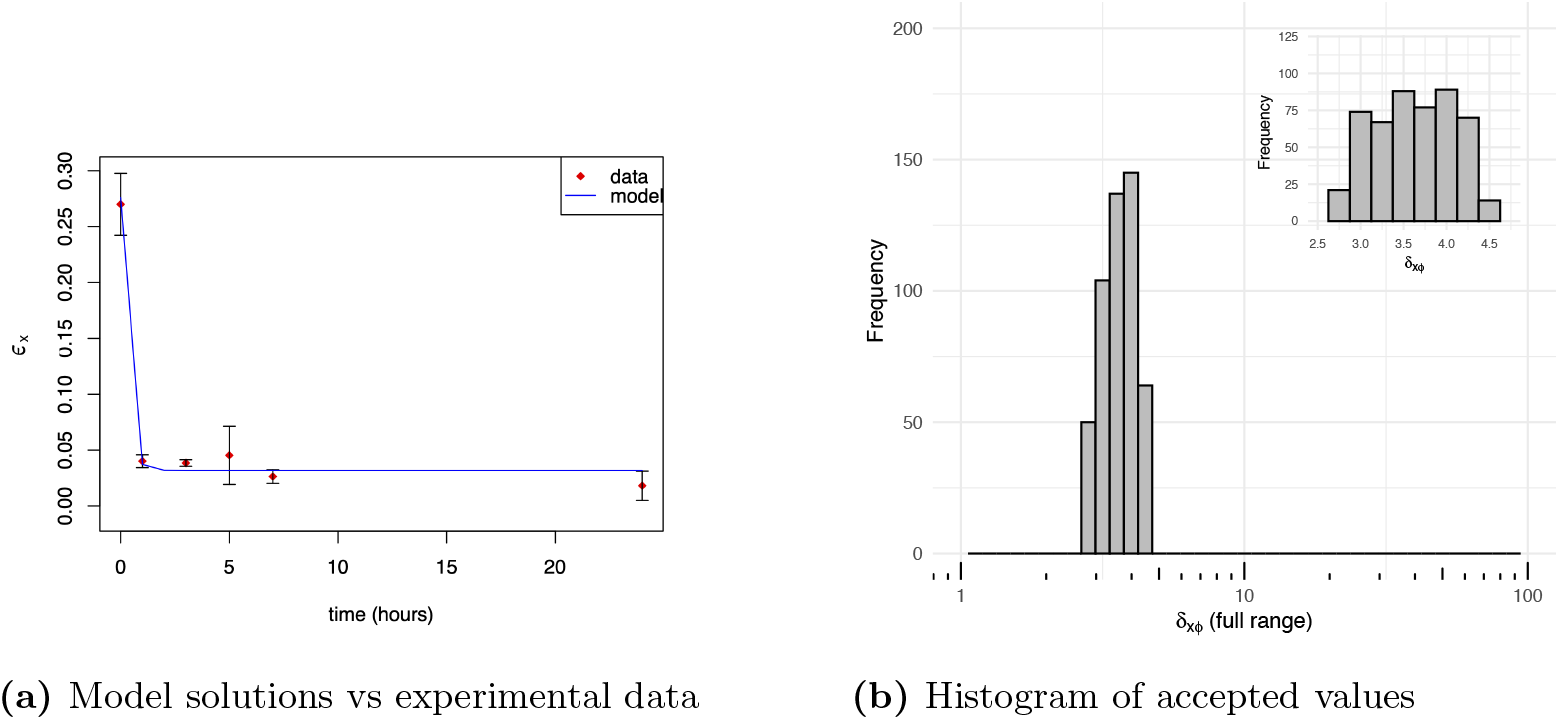
Model solutions to Eq (19). Numerical solutions using the parameter estimate of *δ*_*xϕ*_ = 3.81 hr^−1^ and experimental data from biofilms treated with undoped fibres (Fig 5a). The error bars indicate the standard deviation of the data (n=3). Fig 5b shows the range of accepted values of *δ*_*xϕ*_ on a log scale (x-axis) using the ABC method, where values were sampled from a uniform distribution ranging from 0 to 100 hr^−1^ (inset: range of accepted values on a linear scaling, focusing on a smaller range of the x-axis). *C* & *B* were held fixed for the ABC method, where *C* = 0.24 & *B* = 0.03.

To further explore the full range of possible values that *δ*_*xϕ*_ can take to get an appropriate fit to the experimental data, the ABC method is used where *δ*_*xϕ*_ is uniformly sampled from the interval [0, 100] hr^−1^. For each sampled value of *δ*_*xϕ*_ the absolute error *ρ* between the experimental data and model solutions is computed as:

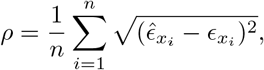

where 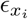 represents the model solution and 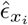 denotes the experimental data. We select 500 samples of *δ*_*xϕ*_ that result in a sufficiently small error (*ρ* ≤ 8.5 × 10^−5^), as errors smaller than this are rarely observed. The distribution of the resulting estimates is shown in Figure 5b. The plot reveals that only a narrow range of values (*δ*_*xϕ*_ ∈ (2.6, 4.6) hr^−1^) could yield model solutions with an acceptable fit to the experimental data (i.e. where *ρ* ≤ 8.5 × 10^−5^), suggesting we can be relatively confident in our default value of *δ*_*xϕ*_ = 3.81.

#### Determining *δ*_*xa*_ **and** *δ*_*ax*_

Next, the parameter estimates for *δ*_*xa*_ and *δ*_*ax*_ (respective death rate of live cells by antimicrobial and uptake rate of antimicrobial by live cells) are determined. Given that the majority of physical disruption occurs within the initial hour of treatment, as evidenced by the experimental findings (Fig 4a), we postulate that subsequent physical disruption is negligible, with cell death primarily driven by antimicrobial ions after the initial hour. Consequently, for the purpose of estimating *δ*_*xa*_, the differential equation representing the change in live cell fraction (*ϵ*_*x*_) with respect to time can be further simplified to:

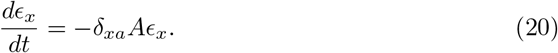

Here, we set the initial conditions *ϵ*_*x*_(1) = *ϵ*_*x*1_ and *A*(1) = *A*_1_ where *ϵ*_*x*1_ and *A*_1_ are given initial values but are also included as fitted parameters. We then focus our analysis on the data for *t* ≥ 1 hr. This approach allows a more targeted examination of the system dynamics. The primary parameters of interest, *δ*_*xa*_ and *δ*_*ax*_, are estimated by fitting the solutions of Eq. (17) and Eq. (20) (the ODEs describing the antimicrobial concentration within the biofilm and the live cell fraction, respectively) to the experimental results from biofilm treatment with silver-doped fibres (red curve in Fig. 4b, with numerical values in Table 4).

The resulting estimates for *δ*_*xa*_ and *δ*_*ax*_ were heavily dependent on the initial guesses that were fed into the algorithm (illustrating an identifiability issue): see Table 5. Furthermore, the estimated value of *δ*_*xa*_ produced by the least squares method maintained the same order of magnitude as the initial guess, where acceptable parameter estimates can be found for *δ*_*xa*_ = *O*(10^−1^) − *O*(10^3^). The relationship between *δ*_*xa*_ and *δ*_*ax*_ is notable. When *δ*_*xa*_ falls within the range of *O*(1) − *O*(10^3^), *δ*_*ax*_ is adjusted to maintain a balanced rate of live cell death, typically around *O*(10^−1^). Conversely, for *δ*_*xa*_ = *O*(10^−1^), *δ*_*ax*_ becomes negligibly small, leading to an underestimation of the live cell death rate. When *δ*_*xa*_ drops below *O*(10^−1^), antimicrobial ions are insufficient to cause significant cell death, even in the absence of ion uptake. Additionally, while estimates for *ϵ*_*x*1_ remain independent of initial guesses, *A*_1_ estimates vary considerably for higher *δ*_*xa*_ values but stabilise around 0.8125 *µ*g ml^−1^ for *δ*_*xa*_ = *O*(10^−1^).

**Table 5.** Parameter estimates for *δ*_*xa*_, *δ*_*ax*_, *ϵ*_*x*1_, *A*_1_ and their associated errors using the least squares method when different initial guesses were used.

| Initial Guess | Estimates |  |  |  | Error |
| --- | --- | --- | --- | --- | --- |
| $(\delta_{\mathbf{x}\mathbf{a}}, \delta_{\mathbf{a}\mathbf{x}})$ | $\delta_{\mathbf{x}\mathbf{a}}$<br>[ml $\mu\text{g}^{-1}$ hr$^{-1}$] | $\delta_{\mathbf{a}\mathbf{x}}$<br>[hr$^{-1}$] | $\epsilon_{\mathbf{x}1}$<br>[-] | $\mathbf{A}_1$<br>[$\mu\text{g}$ ml$^{-1}$] | |
| $(10^{-1}, 10)$ | 0.64 | $9.1 \times 10^{-7}$ | 0.068 | 0.8125 | $2.37 \times 10^{-4}$ |
| $(10^{-1}, 0.1)$ | 0.64 | $2.6 \times 10^{-5}$ | 0.068 | 0.8125 | $2.37 \times 10^{-4}$ |
| $(10^{-1}, 10^{-10})$ | 0.64 | $3.5 \times 10^{-5}$ | 0.068 | 0.8125 | $2.37 \times 10^{-4}$ |
| $(1, 10)$ | 1 | 24 | 0.068 | 0.78 | $1.15 \times 10^{-3}$ |
| $(10^1, 10)$ | 13.54 | 320 | 0.068 | $1.3 \times 10^{-5}$ | $4.04 \times 10^{-4}$ |
| $(10^2, 10)$ | $10^2$ | $3.5 \times 10^3$ | 0.068 | 0.089 | $2.38 \times 10^{-3}$ |
| $(10^3, 10)$ | $10^3$ | $3.3 \times 10^4$ | 0.068 | $3.9 \times 10^{-7}$ | $2.26 \times 10^{-3}$ |

To address this identifiability issue, for subsequent solutions, two parameter sets are chosen (see Table 6); the parameter set, Θ_1_, is chosen so that *δ*_*xa*_*/δ*_*ax*_ = *O*(10^−1^) and Θ_2_ is chosen where *δ*_*ax*_ must be negligibly small.

**Table 6.** Parameter estimates for *δ*_*xa*_, *δ*_*ax*_, *A*_1_ **used for model solutions**.

| Parameter | $\Theta_1$ | $\Theta_2$ |
| --- | --- | --- |
| $\delta_{\mathbf{x}\mathbf{a}}$ [ml $\mu\text{g}^{-1}$ hr$^{-1}$] | 13.54 | 0.64 |
| $\delta_{\mathbf{a}\mathbf{x}}$ [hr$^{-1}$] | $3.2 \times 10^2$ | $9.1 \times 10^{-7}$ |
| $\mathbf{A}(\mathbf{1}) = \mathbf{A}_1$ [$\mu\text{g}$ ml$^{-1}$] | $1.3 \times 10^{-5}$ | 0.8125 |
| $\epsilon_{\mathbf{x}}(\mathbf{1}) = \epsilon_{\mathbf{x}1}$ [-] | 0.068 | 0.068 |

The solutions to (17) & (20) using the two parameter sets are shown in Fig 6, where good agreement can be seen for either parameter set between the solution to (20) and the data, and different behaviour between the two sets for (17). Using the Θ_1_ dataset, the antimicrobial concentration initially increases slowly but then increases at a faster rate as the live cell fraction decreases over time, resulting in less uptake of the ions (Fig 6c). This increase lasts until *t* = 8 hrs, at which point ions are no longer released and the live cell fraction is negligible, thus resulting in no more uptake of antimicrobial. Conversely, using the Θ_2_ dataset, there is a linear increase in the ion concentration until *t* = 8 hrs, at this point the concentration remains constant (Fig 6d). This is because the estimated value for *δ*_*ax*_ is negligibly small so there is effectively no uptake of antimicrobial, resulting in a higher constant concentration of 6.5 *µ*g ml^−1^ at *t* = 8 hrs compared to solutions using the Θ_1_ dataset.

**Fig 6.**
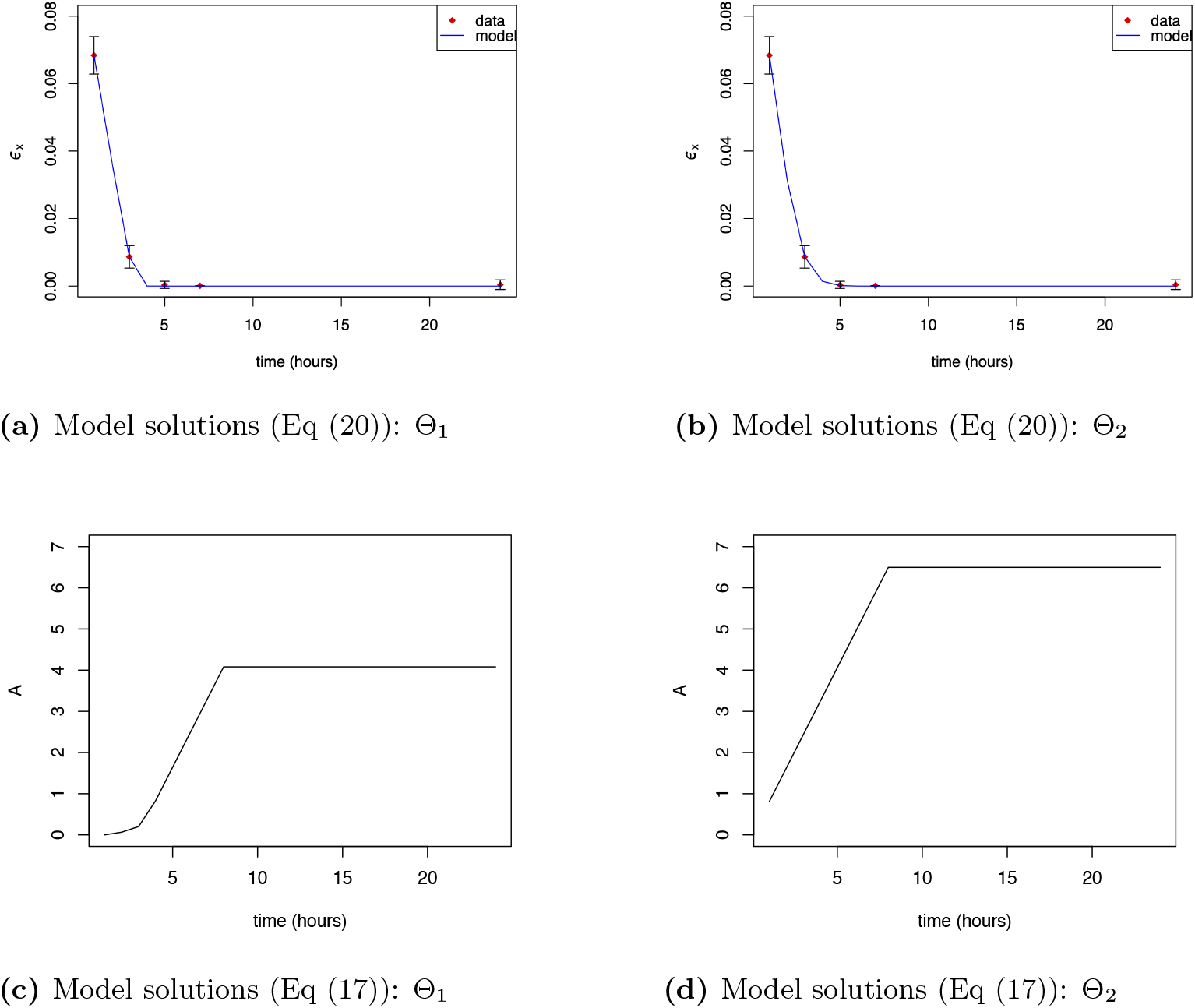
Model solutions to Eqs (20) & (17). Solutions using the Θ_1_ parameter estimates, *δ*_*xa*_=13.54 ml *µ*g^−1^ hr^−1^ and *δ*_*ax*_ = 3.2 × 10^2^ hr^−1^, or the Θ_2_ parameter estimates, *δ*_*xa*_=0.64 ml *µ*g^−1^ hr^−1^ and *δ*_*ax*_ = 9.1 × 10^−7^ hr^−1^. In Figs 6a & 6b, model solutions are compared to experimental data, where the error bars indicate the standard deviation of the data (n=3).

Next, the ABC method is applied, where parameter values are accepted if the error (*ρ*) is less than 1 × 10^−5^ between model solutions and experimental data. Initially, *δ*_*ax*_ (and *A*_1_) are assigned to their Θ_1,2_ parameter sets, while *δ*_*xa*_ values are sampled uniformly from the interval [0, 2 × 10^3^] ml *µ*g^−1^ hr^−1^. The accepted *δ*_*xa*_ values yielding a good fit fall within a much narrower range than the sampling distribution (see Figs 7a and 7b), where,

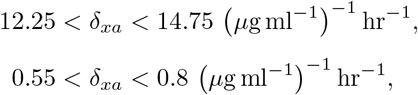

for Θ_1_ and Θ_2_ respectively. This therefore informs us of our confidence in *δ*_*xa*_ given that *δ*_*ax*_ is fixed.

**Fig 7.**
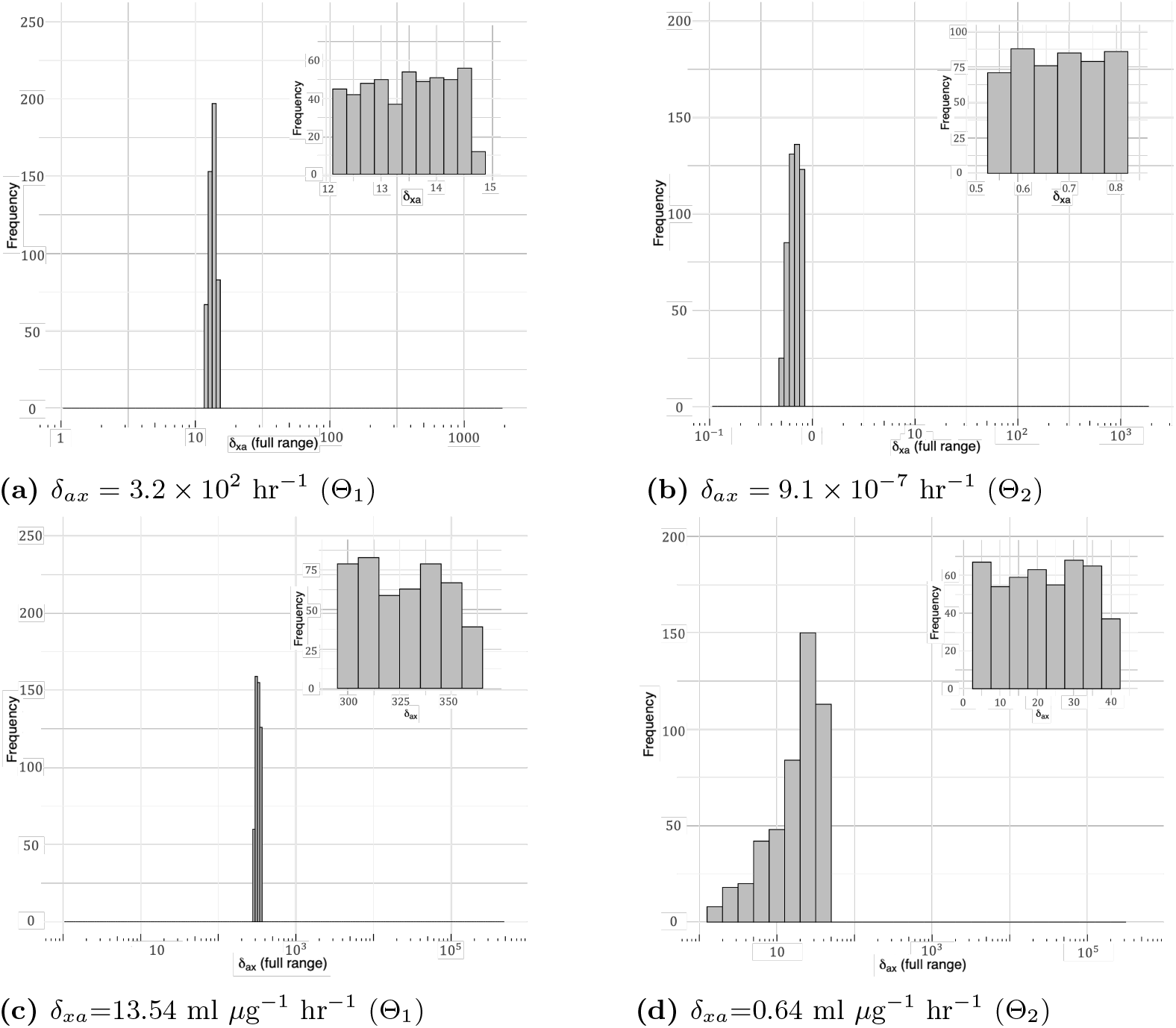
Estimated parameter distributions. The distribution ranges of accepted values of *δ*_*xa*_ and *δ*_*ax*_ that produced minimal error between model solutions and experimental data on a log scale (x-axis) using the ABC method. The inset plot displays the range of accepted values on a linear scale, focusing on a smaller range of the x-axis.

Next, *δ*_*xa*_ (and *A*_1_) are fixed using the Θ_1,2_ parameter sets, while *δ*_*ax*_ values are sampled uniformly from the interval [0, 5 × 10^5^] hr^−1^. Once again, for both parameter sets, the ranges of accepted parameter values were much smaller than the range from which the values were sampled on (see Figs 7c and 7d), where,

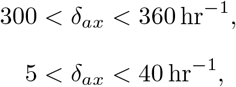

for Θ_1_ and Θ_2_ respectively. However it is noted that when *δ*_*xa*_ is from Θ_2_, the ABC method encounters difficulty in finding suitable *δ*_*ax*_ values, necessitating an increased error tolerance to *ρ* < 1 × 10^−2^. Hence, parameter sets from Θ_2_ exhibit poorer fit.

Finally, using the Θ_1,2_ parameter sets, we used the ABC method to simultaneously sample *δ*_*xa*_ and *δ*_*ax*_ values while fixing only *A*_1_ (Fig 8). For both parameter sets, *δ*_*xa*_ values span the entire range, with a higher frequency around *δ*_*xa*_ = *O*(10^3^). Conversely, *δ*_*ax*_ values cover a narrower range.

**Fig 8.**
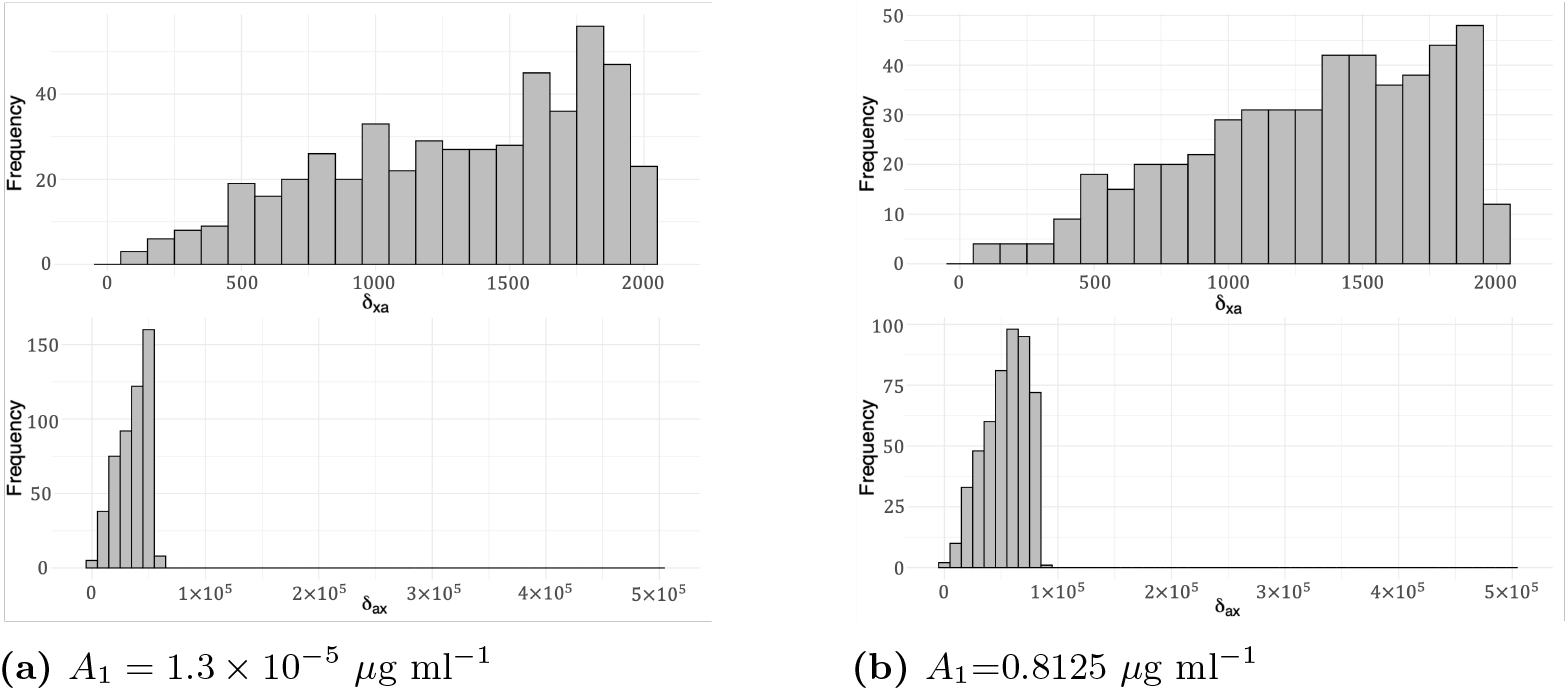
Estimated parameter distributions. The distribution ranges of accepted values of *δ*_*xa*_ and *δ*_*ax*_ that produced minimal error between model solutions and experimental data using the 2 parameter cases for *A*_1_ given in Table 6.

The ABC method confirms the coupling between *δ*_*xa*_ and *δ*_*ax*_. Fixing one parameter restricts the possible values of the other parameter to a narrow range for a good fit. However, sampling both parameters results in a broad range of possible values. Similar trends are observed with the least squares method: larger *δ*_*xa*_ values correlate with *δ*_*xa*_*/δ*_*a*_ = *O*(10^−1^) ml *µ*g^−1^, while smaller *δ*_*xa*_ values necessitate negligibly small *δ*_*ax*_. Decoupling these parameters would require additional experimental data, such as measuring ion concentration within the biofilm during treatment. In subsequent sections, we initially employ both the Θ_1,2_ values for *δ*_*ax*_ and *δ*_*xa*_, comparing solutions obtained from the full PDE model to experimental data, demonstrating that Θ_1_ yeilds more realistic results when considering biofilm eradication time.

### Model solutions using estimated parameters

In the thin biofilm case, *O*(*z, t*) ∼ 1 throughout, and assuming *γ*_*ox*_, *γ*_*ax*_ ≪ 1, we have 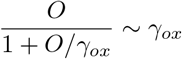 and 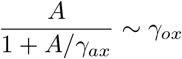, reducing the system (1), (2) and (4)-(6) to:

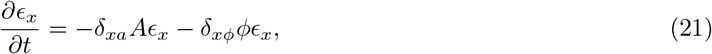

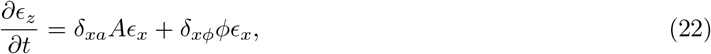

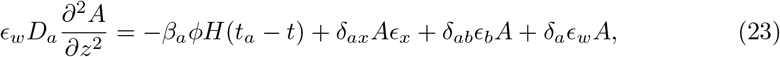

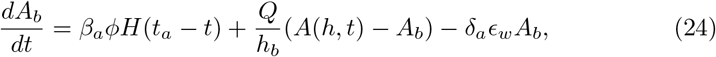

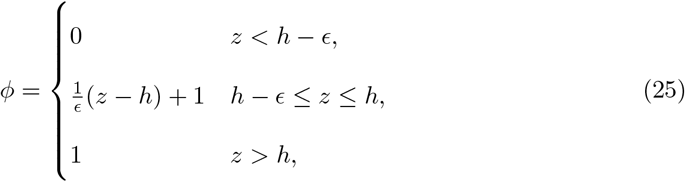

recalling the redefinition of *γ*_*ox*_*δ*_*xa*_ ↦ *δ*_*xa*_ and *γ*_*ax*_*δ*_*ax*_ ↦ *δ*_*ax*_, where *δ*_*xa*_ and *δ*_*ax*_ henceforth denote the rescaled parameters. The model is subject to the boundary and initial conditions (10)-(12), oxygen being decoupled from the system by these assumptions. The two possible parameter sets for *δ*_*xa*_ and *δ*_*ax*_ given in Table 6 are initially both considered.

Unless specified otherwise, the base case values for all other parameters are in Table 7, alongside the justification for each parameter value. More detailed explanations for the parameters can be found in [34].

**Table 7.**
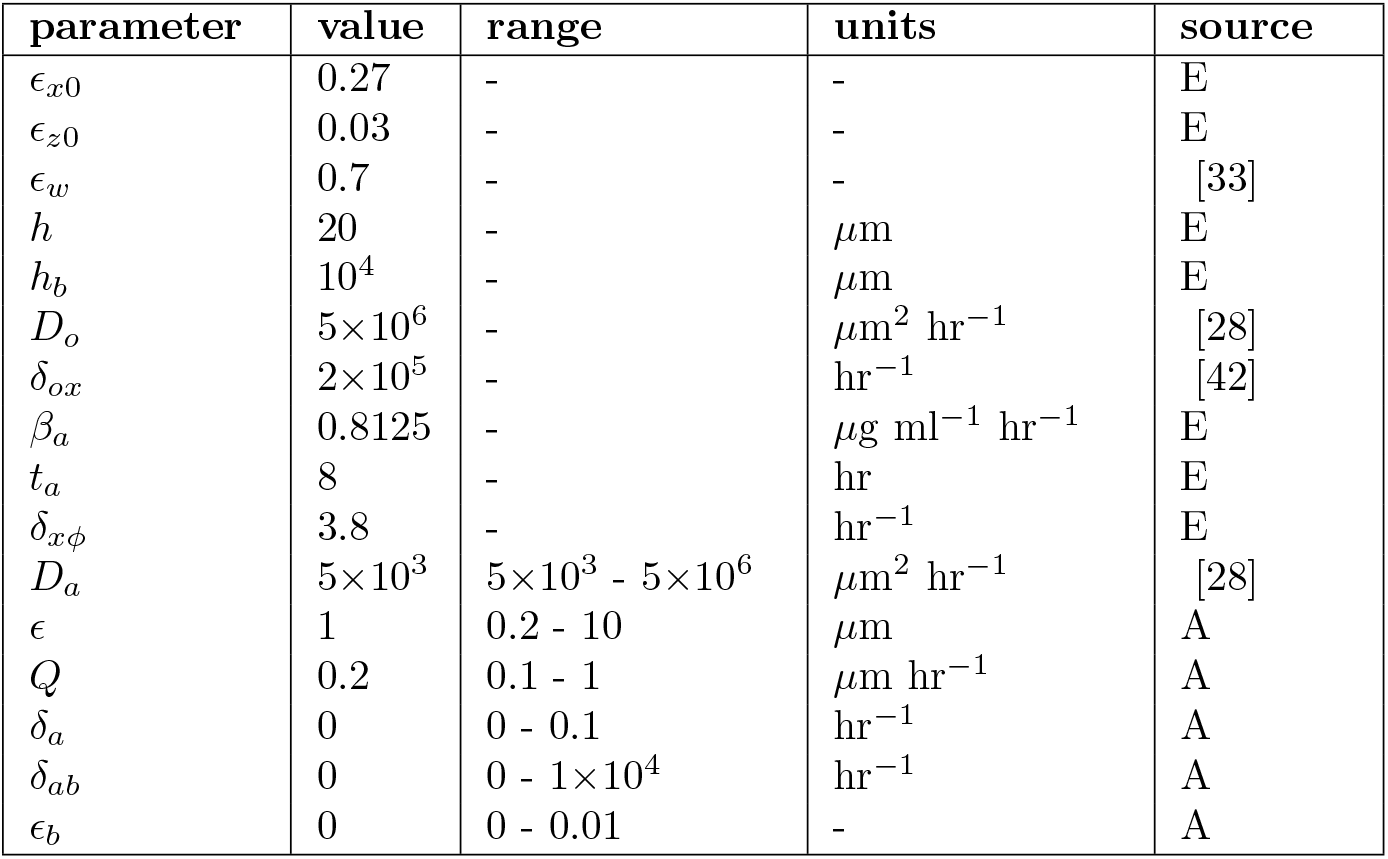
Base case parameter values for Eqs (1)-(6), excluding *δ*_*xa*_ and *δ*_*ax*_, which can be found in Table 6. Sources for parameter values are provided: values are either determined from literature, estimated from data (E), or assumed to give qualitatively desirable results (A). Where appropriate, the ranges for parameters that are varied and literature citations are provided.

| parameter | value | range | units | source |
| --- | --- | --- | --- | --- |
| $\epsilon_{x0}$ | 0.27 | - | - | E |
| $\epsilon_{z0}$ | 0.03 | - | - | E |
| $\epsilon_w$ | 0.7 | - | - | [33] |
| $h$ | 20 | - | $\mu\text{m}$ | E |
| $h_b$ | $10^4$ | - | $\mu\text{m}$ | E |
| $D_o$ | $5 \times 10^6$ | - | $\mu\text{m}^2 \text{ hr}^{-1}$ | [28] |
| $\delta_{ox}$ | $2 \times 10^5$ | - | $\text{hr}^{-1}$ | [42] |
| $\beta_a$ | 0.8125 | - | $\mu\text{g ml}^{-1} \text{ hr}^{-1}$ | E |
| $t_a$ | 8 | - | hr | E |
| $\delta_{x\phi}$ | 3.8 | - | $\text{hr}^{-1}$ | E |
| $D_a$ | $5 \times 10^3$ | $5 \times 10^3 - 5 \times 10^6$ | $\mu\text{m}^2 \text{ hr}^{-1}$ | [28] |
| $\epsilon$ | 1 | 0.2 - 10 | $\mu\text{m}$ | A |
| $Q$ | 0.2 | 0.1 - 1 | $\mu\text{m hr}^{-1}$ | A |
| $\delta_a$ | 0 | 0 - 0.1 | $\text{hr}^{-1}$ | A |
| $\delta_{ab}$ | 0 | 0 - $1 \times 10^4$ | $\text{hr}^{-1}$ | A |
| $\epsilon_b$ | 0 | 0 - 0.01 | - | A |

#### Testing parameter regimes: comparing model solutions for Θ_1_ **and** Θ_2_

We begin by evaluating which parameter set, Θ_1_ or Θ_2_, is more biologically realistic. Figs. 9 and 10 present solutions to Eqs. (21)-(25) using the base case parameter values from Table 7, along with the parameter sets Θ_1_ and Θ_2_ for *δ*_*xa*_ and *δ*_*ax*_, as listed in Table 6. In all subsequent figures, the biofilm height is fixed at *h* = 20, with *z* > *h* representing the bulk fluid.

**Fig 9.**
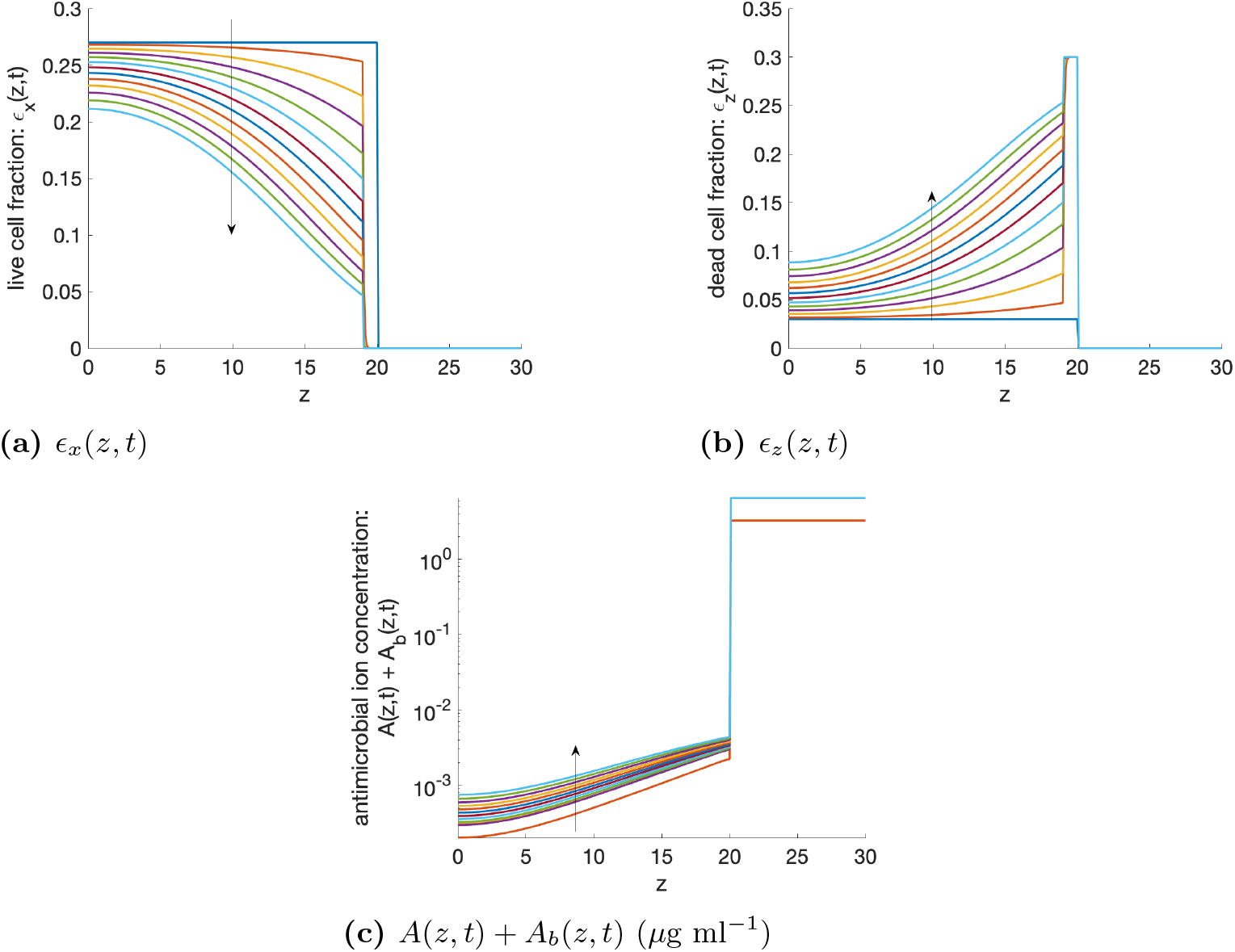
Numerical solutions for (21)-(24). Solutions are in steps of *t* = 4 using parameters given in Table 7 and Θ_1_ parameter values for *δ*_*xa*_ and *δ*_*ax*_ in Table 6. Here Θ_1_ represents a regime in which a higher antimicrobial-induced death rate is balanced by a higher antimicrobial uptake rate. Arrows represent the direction of the change in variables in increasing time.

**Fig 10.**
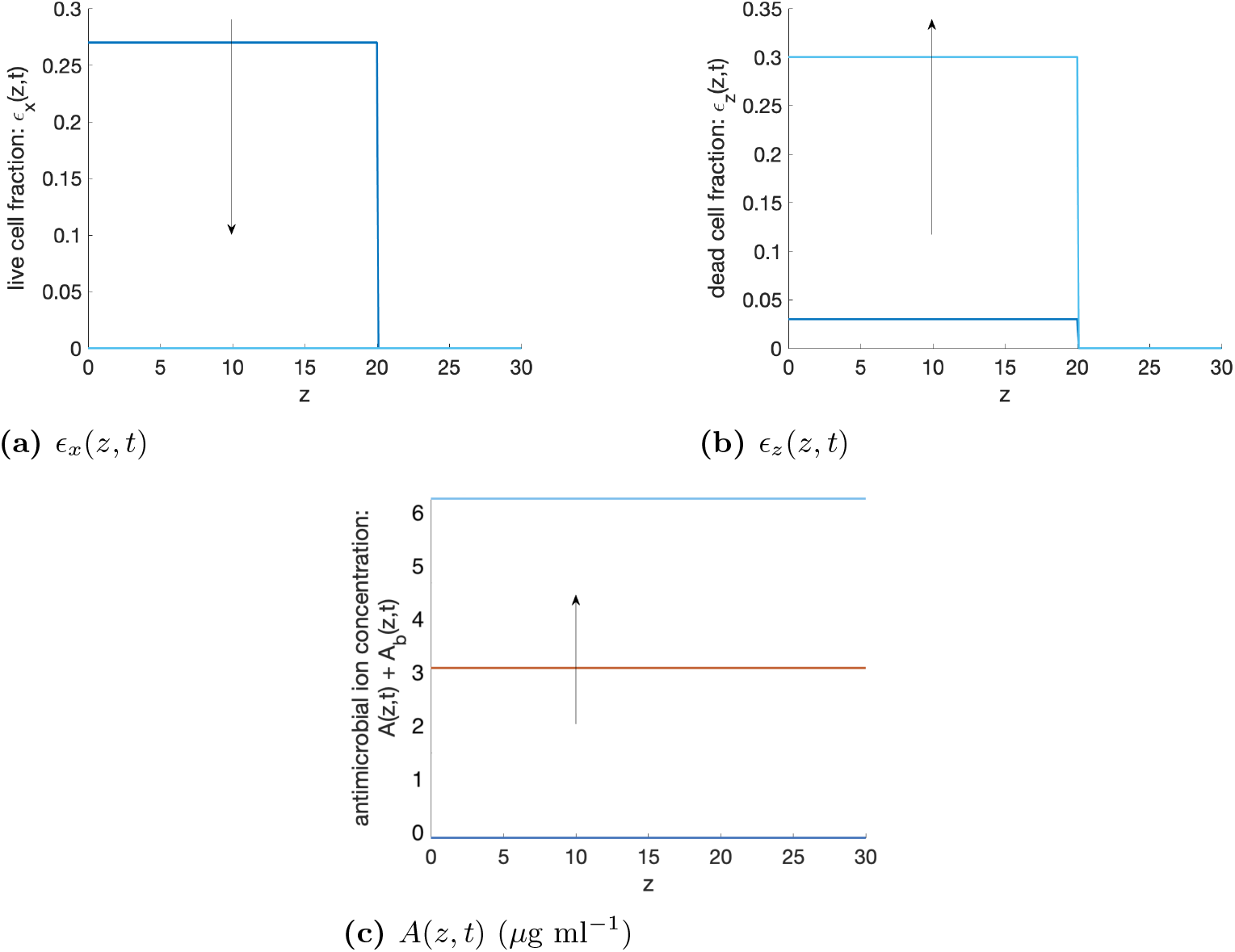
Numerical solutions for (21)-(24). Solutions are in steps of *t* = 4 using parameters given in Table 7 and Θ_2_ parameter values for *δ*_*xa*_ and *δ*_*ax*_ in Table 6. Here, Θ_2_ represents a regime in which a lower antimicrobial-induced death rate is accompanied by negligible antimicrobial uptake. Arrows represent the direction of the change in variables in increasing time.

Using parameter set Θ_1_, after 48 hrs of treatment, the fraction of live cells decreases over time, but is not eradicated (Fig 9a). However, running the simulations for longer, the live cells are eradicated eventually (after approximately 100 hrs, results not shown). We note that the cells in the top 1 *µ*m of the biofilm are eradicated due to physical disruption caused by the BG fibres (Fig 9b). The antimicrobial ion concentration within the biofilm increases with time as ions are released from the fibres and ions diffuse in from the biofilm/bulk liquid interface as the ion concentration in the bulk liquid is much higher than within the biofilm (Fig 9c).

Solutions using the Θ_2_ parameter values are qualitatively different to solutions using the Θ_1_ parameter values (Fig 10), where in this case, bacteria are eradicated quickly (Figs 10a & 10b) due to a negligible uptake of antimicrobial ions by bacteria (recall that *δ*_*ax*_ is very small in the Θ_2_ parameter set). Thus, there is a much higher concentration of antimicrobial ions within the biofilm, where concentrations within the biofilm are the same as concentrations outside the biofilm (Figs 10c).

From experimental work, biofilms treated with silver-doped fibres are not fully eradicated by 24hrs (see Fig 4), therefore the Θ_1_ parameter set is more realistic. Hence, in all future solutions, the Θ_1_ parameter set is used as the base case parameters.

In the following sections we investigate the effect of the parameters that we have been unable to estimate from the data, namely *ϵ, Q* and *D*_*a*_, on the treatment efficacy of BG fibres against biofilms.

#### Investigating the effect of *ϵ* on treatment efficacy

The depth of biofilm penetration by the fibres is captured by *ϵ*. Decreasing *ϵ* from its base case value of *ϵ* = 1 to *ϵ* = 0.2 *µ*m results in an increase in the fraction of remaining live cells (*ϵ*_*x*_) after 48 hours of treatment (see Fig 11a) and also results in lower concentrations of antimicrobial (*A*) within the biofilm as fewer ions are released directly into the biofilm area (see Fig 11b). Conversely, increasing *ϵ* decreases the fraction of live cells after 48 hours of treatment, where total live cell eradication occurs by 48 hours when *ϵ* = 10 *µ*m. This is due to fibres penetrating deeper within the biofilm, resulting in more cells dying through physical disruption and also facilitates a higher concentration of antimicrobial ions within the biofilm.

**Fig 11.**
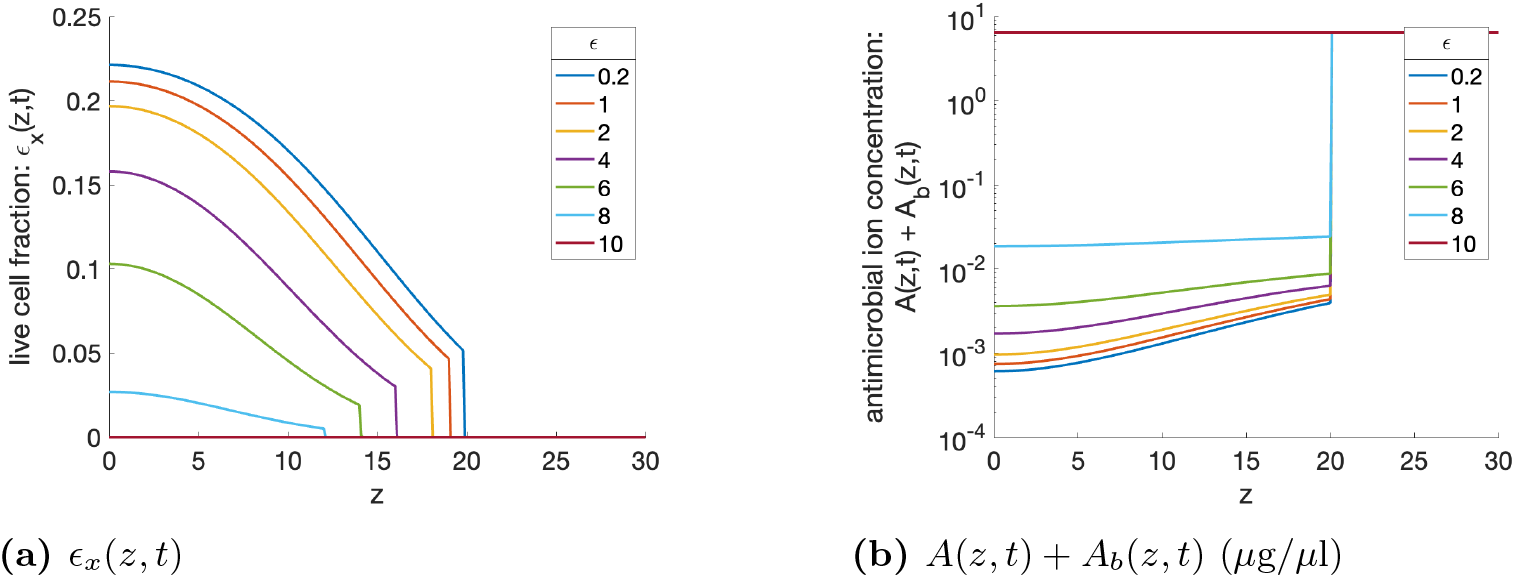
Numerical solutions for (21)-(24). Solutions are evaluated at *t* = 48 hours using parameters given in Table 7 and the Θ_1_ parameter set for *δ*_*xa*_ and *δ*_*ax*_, for different values of *ϵ*.

#### Investigating the effect of *Q* on treatment efficacy

The flux of ions between the biofilm/bulk liquid interface is captured by the parameter *Q*. Increasing *Q* from its base value of *Q* = 0.2 *µ*m hr^−1^ results in ions in the bulk liquid travelling into the biofilm much faster, therefore leading to higher ion concentrations within the biofilm, resulting in faster death of live cells (Figs 12a & 12b), where live cell eradication occurs by 48 hours when *Q* = 1 *µ*m hr^−1^. Reducing the flux to *Q* = 0.1 *µ*m hr^−1^ has the opposite effect.

**Fig 12.**
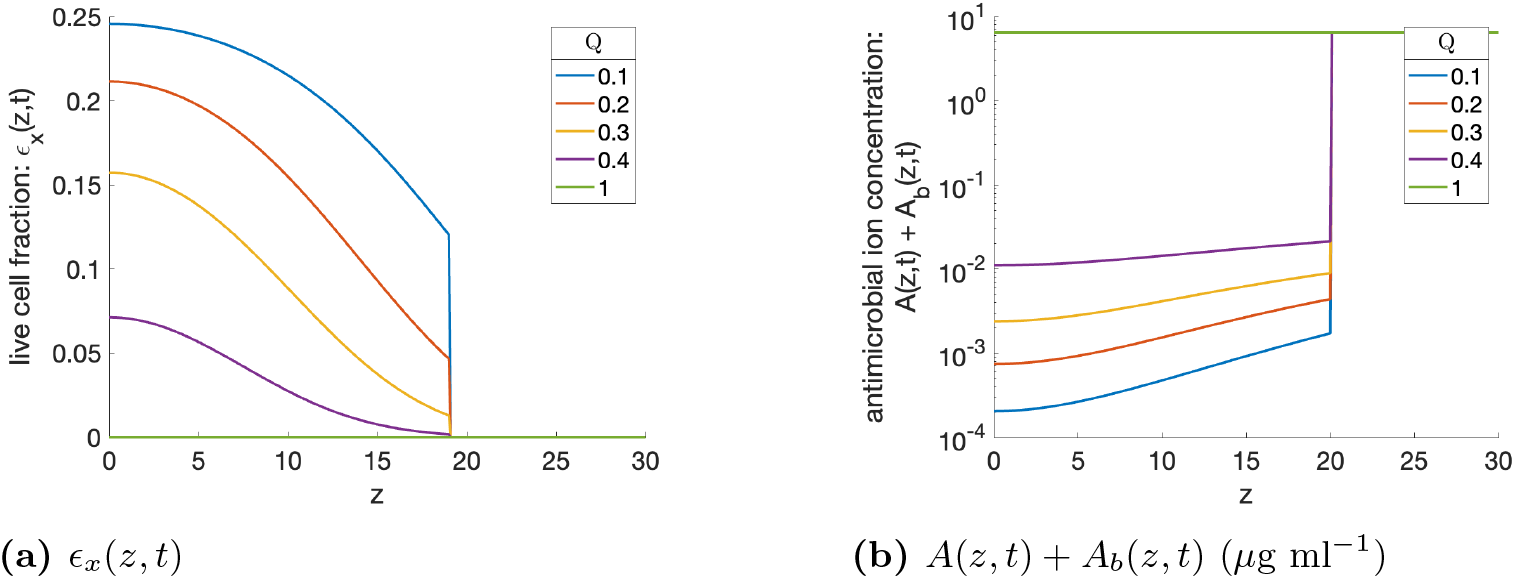
Numerical solutions for (21)-(24). Solutions are evaluated at *t* = 48 hours using parameters given in Table 7 and the Θ_1_ parameter set for *δ*_*xa*_ and *δ*_*ax*_, for different values of *Q*.

#### Investigating the effect of *D*_*a*_ on treatment efficacy

The value of *D*_*a*_ (diffusion coefficient of antimicrobial ions) differs depending on whether the silver is released as ions or in nanoparticle form. Ions, being smaller than nanoparticles, have a larger diffusion coefficient (where 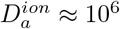 and 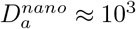). Here we investigate the effect of increasing *D*_*a*_, to see the effect this has on biofilm treatment. For the smaller diffusion coefficient, the antimicrobial concentration decreases with increasing depth into the biofilm (Fig 13b). Conversely, increasing the diffusion coefficient results in a uniform concentration of *A* within the biofilm. Consequently, with large *D*_*a*_ there is uniform death of live cells throughout the biofilm (Fig 13a), whereas more cells are killed at the top of the biofilm compared to the bottom when *D*_*a*_ is smaller. Importantly, despite the antimicrobial penetrating to differing depths with changes to *D*_*a*_, biofilm eradication occurs within a similar timeframe. Specifically, increasing antimicrobial diffusion within the biofilm does not appear to enhance the likelihood or speed of eradication. Here, at low diffusion, the antimicrobial reaches high local concentrations near the biofilm surface, causing substantial cell death that progressively spreads towards the substratum as surface cells die, whereas at high diffusion the antimicrobial is distributed more uniformly throughout the biofilm, producing a less intense but more widespread attack. Consequently, these competing effects result in similar overall eradication timescales.

**Fig 13.**
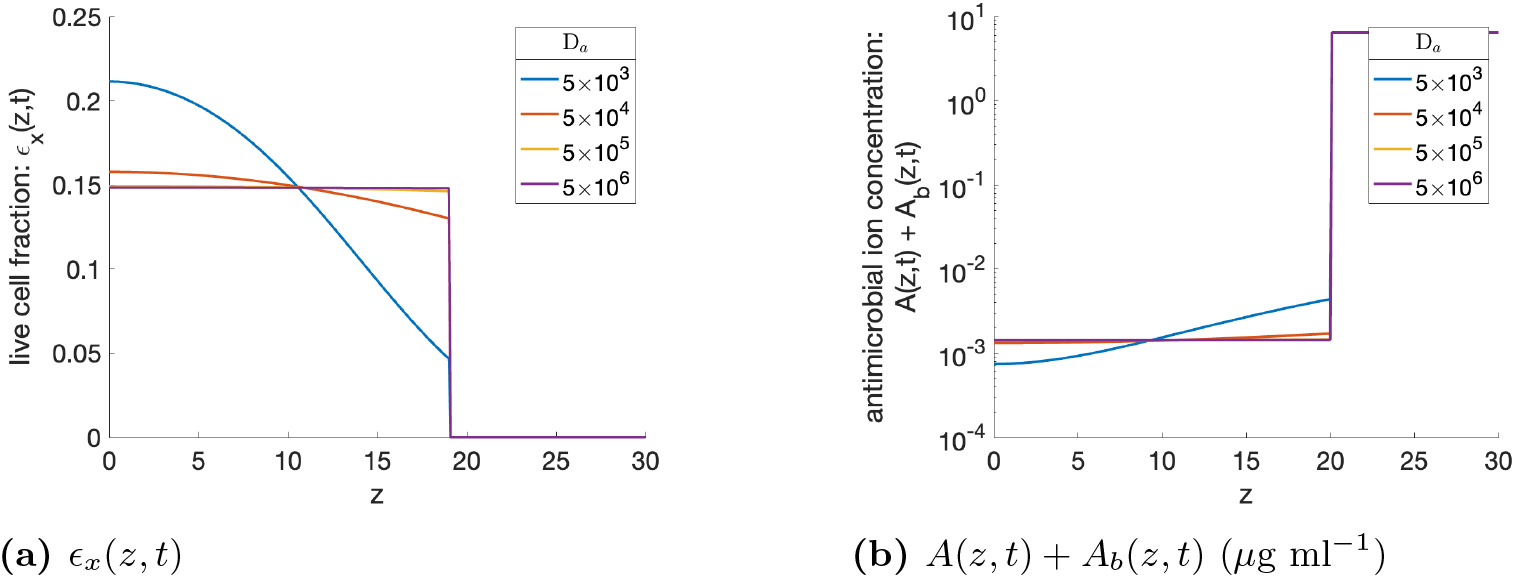
Numerical solutions for (21)-(24). Solutions are evaluated at *t* = 48 hours using parameters given in Table 7 and the Θ_1_ parameter set for *δ*_*xa*_ and *δ*_*ax*_, for different values of *D*_*a*_.

**Fig 14.**
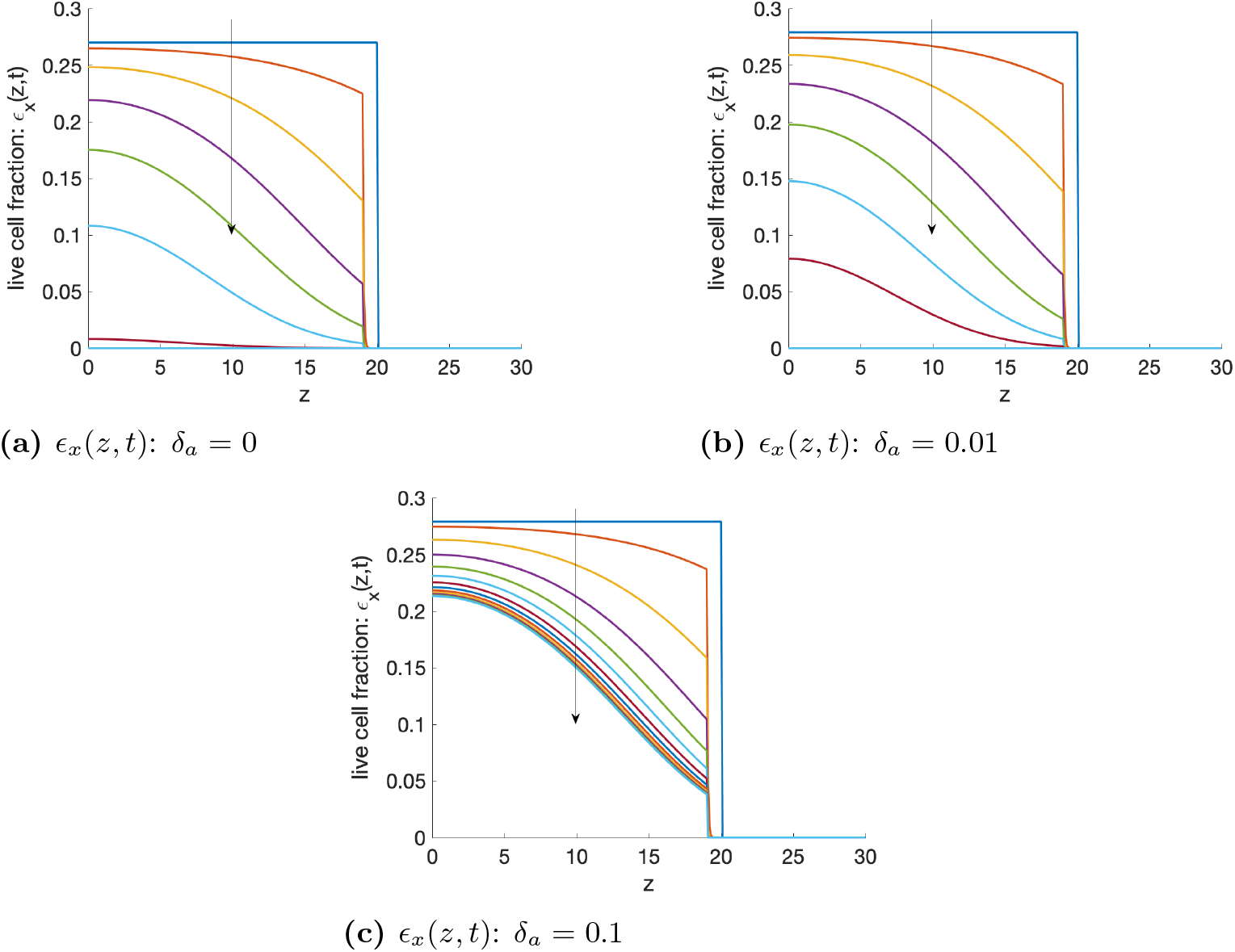
Numerical solutions for (21). Solutions are in steps of *t* = 4 using parameters given in Table 7, *δ*_*ab*_ = 0 and the Θ_1_ parameter set for *δ*_*xa*_ and *δ*_*ax*_, where either *δ*_*a*_ = 0 (Fig 14a), *δ*_*a*_ = 0.01 (Fig 14b) or *δ*_*a*_ = 0.1 (Fig 14c). Arrows represent the direction of the change in variables in increasing time.

### Modelling mechanisms for removal of antimicrobial ions

We now explore two potential mechanisms that could diminish the antimicrobial effectiveness of silver. Firstly, experimental findings from [17] indicate that silver ions exhibit instability in typical cell culture media, reacting with sodium chloride to form silver chloride, which, unlike silver, lacks antimicrobial properties. This process effectively removes antimicrobial silver from the system (modelled by the parameter *δ*_*a*_), contributing to decreased antimicrobial activity. Secondly, the binding of positively charged silver ions to negatively charged EPS, reducing the availability of free ions, and hence antimicrobial efficacy (modelled by the parameter *δ*_*ab*_).

We now incorporate the presence of EPS (*ϵ*_*b*_) into the model and assume it comprises approximately 1% of the biofilm mass [34], i.e. *ϵ*_*b*_ = 0.01 (increased from *ϵ*_*b*_=0 in the previous section). Using the same initial conditions for *ϵ*_*x*0_ and *ϵ*_*z*0_ as listed in Table 7, and applying the conservation of volume equation (7), we obtain *ϵ*_*w*_ = 0.69. For model simulations, we use the parameter set Θ_1_, previously shown to provide more realistic predictions. Here, we apply a higher exchange rate of antimicrobial ions, *Q* = 1 *µ*m hr^−1^, at the biofilm–bulk liquid interface to model a scenario in which complete biofilm eradication occurs after approximately 48 hours. This allows us to investigate how the loss mechanisms governed by *δ*_*a*_ and *δ*_*ab*_ may inhibit or delay biofilm eradication under otherwise effective treatment conditions.

We begin by considering each new mechanism in turn. Initially, we set *δ*_*ab*_ = 0 (indicating EPS does not remove antimicrobial ions by binding to them) and consider *δ*_*a*_ = 0.01 hr^−1^ (indicating antimicrobial silver ions react with sodium chloride to form non-antimicrobial silver chloride). Here, live cells are still eradicated, but at a slower rate than when *δ*_*a*_ = 0 (14b). However when *δ*_*a*_ is increased tenfold, the biofilm is no longer eradicated within 48 hrs (14c) or after extended periods of time (e.g. 96 hours, data not shown).

This unsuccessful eradication of bacteria is due to the antimicrobial concentration dynamics. In the first 8hrs the antimicrobial concentration increases due to the burst release of ions from the fibres (Fig 15a). However, after 8hrs, ions are no longer released from fibres and the concentration within the biofilm begins to decrease as the ions are removed from the system at a fast rate (Fig 15b). Finally, the ion concentration tends towards zero (Fig 15c), therefore the live cells are no longer killed and reach a steady state.

**Fig 15.**
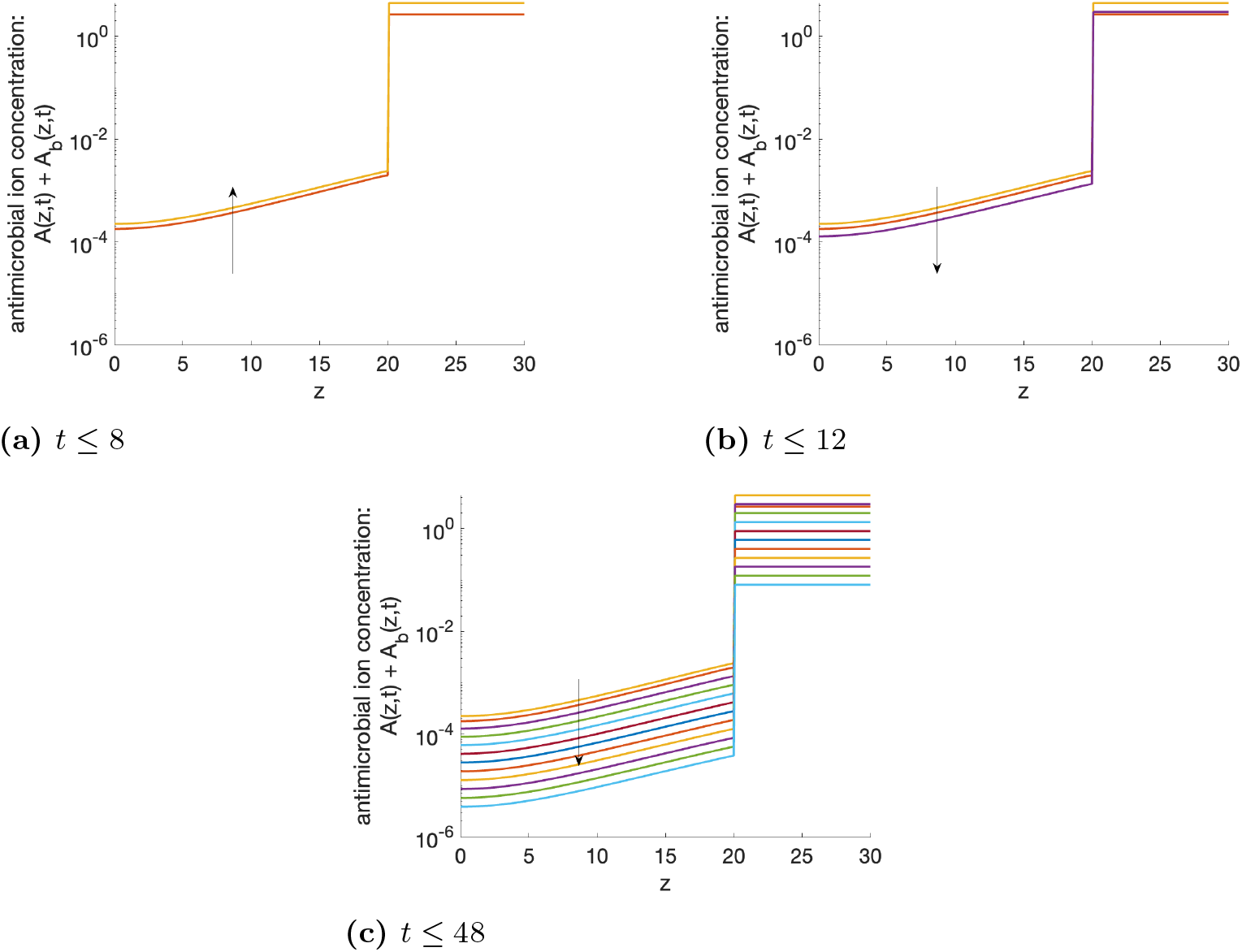
Numerical solutions for (4), (5). Solutions are in steps of *t* = 4 using parameters given in Table 7, *δ*_*ab*_ = 0 and the Θ_1_ parameter set for *δ*_*xa*_ and *δ*_*ax*_, when *δ*_*a*_ = 0.1. Arrows represent the direction of the change in variables in increasing time.

Setting *δ*_*a*_ = 0 hr^−1^ (i.e. there is no removal of ions through a sink term) and looking at the effect of *δ*_*ab*_ ≠ 0 hr^−1^ (i.e. antimicrobial ion removal through binding with EPS), yielded similar qualitative behaviours, although in this case, it is required that *δ*_*ab*_ ≳ 5 × 10^3^ hr^−1^, to have any substantial effect as *ϵ*_*b*_ is very small (Fig 16a). However, the key difference in this ion removal mechanism is that biofilm eradication is always predicted to occur in the long term. This is because ions in the bulk liquid are not removed by EPS, since none is present in that compartment, resulting in a steady flux of antimicrobial ions into the biofilm from the bulk liquid 16b). In contrast, when *δ*_*a*_ ≠ 0 hr^−1^ and *δ*_*ab*_ = 0 hr^−1^ silver ions are removed from both the biofilm and the bulk liquid due to their reaction with sodium chloride, leading to a reduced overall concentration and flux into the biofilm.

**Fig 16.**
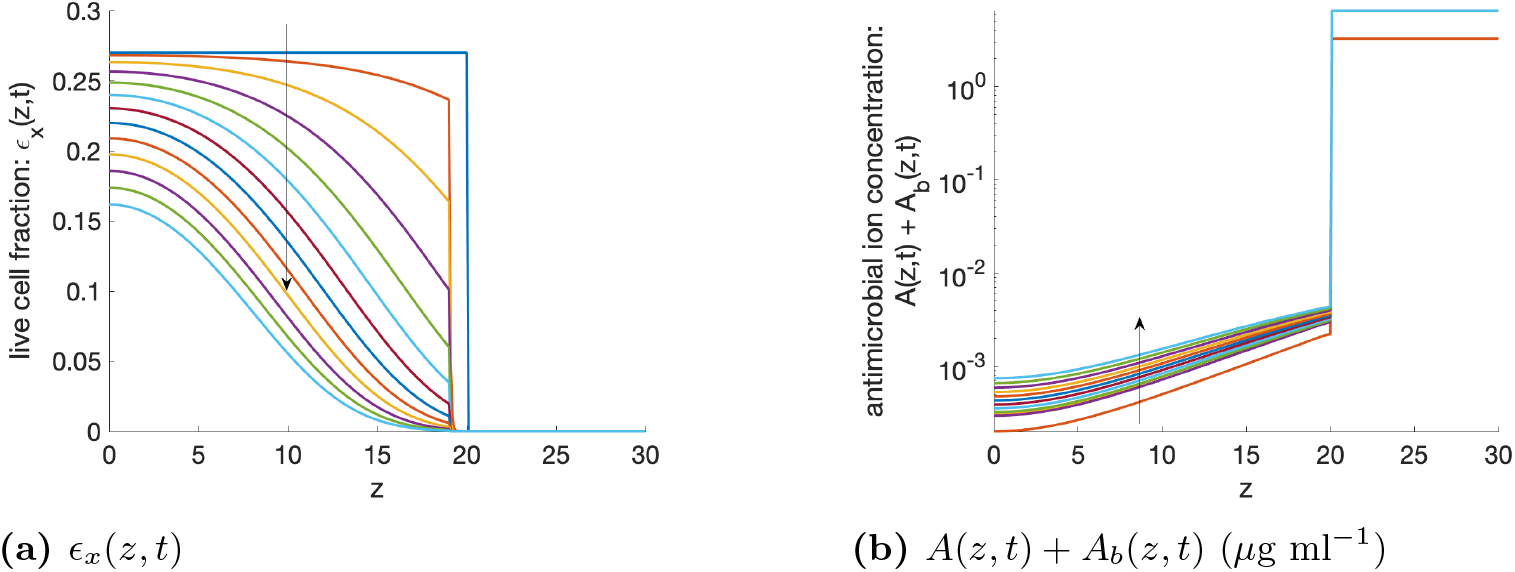
Numerical solutions for (21) and (4), (5). Solutions are in steps of *t* = 4 using parameters given in Table 7, *δ*_*a*_ = 0 and the Θ_1_ parameter set for *δ*_*xa*_ and *δ*_*ax*_, when *δ*_*ab*_ = 5 × 10^3^. Arrows represent the direction of the change in variables in increasing time.

Antimicrobial ion removal from the system, whether through natural sink processes or EPS binding, delays or inhibits biofilm eradication. This effect is compounded when both removal mechanisms are present simultaneously (i.e. *δ*_*a*_ ≠ 0, *δ*_*ab*_ ≠ 0), underscoring the need to optimise BG fibre formulations to minimise these removal pathways.

## Conclusion

In this study, we have developed a non-linear PDE model to assess the efficacy of antimicrobial treatment using BG fibres doped with antimicrobial ions against established biofilm infections. Our model focused on a dynamic equilibrium within the biofilm, initially excluding EPS, with constant water proportion and fixed biofilm height.

The model was parameterised from experimental data to estimate key parameters such as the death rate of bacteria due to antimicrobial ions and physical disruption, as well as the uptake rate of antimicrobial ions by bacteria. By fitting two sets of parameters to experimental data, one with a significant antimicrobial uptake rate and another with negligible uptake, we identified that the set with significant uptake provided more realistic predictions. This demonstrates the importance of accurately capturing antimicrobial interactions within the biofilm for predictive modelling.

An intriguing finding from our simulations was that the duration required for biofilm eradication remained relatively consistent regardless of the antimicrobial diffusion rate. This suggests that factors beyond diffusion kinetics, such as microbial growth dynamics and biofilm architecture, play crucial roles in determining treatment efficacy. Thus, the model predictions have enabled us to assess the relative importance of key processes, indicating that for thin biofilms diffusion is not the limiting factor in these scenarios and that biological mechanisms likely have a more dominant influence on treatment outcomes.

Next we incorporated a sink term for antimicrobial ions in the media, based on previous experimental work [17]. This adjustment improved the model’s ability to predict non-zero steady-state live cell fractions, thereby enhancing its representativeness of real-world scenarios. Additionally, by modelling EPS binding to antimicrobial ions while maintaining a constant EPS fraction, we observed similar qualitative results. Our results shows that treatment efficacy is highly dependent on antimicrobial removal rates, which is crucial to consider in practice. In particular, if removal rates are too high, treatment efficacy diminishes significantly. As EPS-mediated removal is a mechanism that can be actively targeted through material design or treatment strategies, optimising BG fibre development to mitigate this process is crucial for improving therapeutic outcomes.

BG fibres present a viable treatment option for biofilms due to their dual mechanism of action. They can cause physical disruption through the mechanical presence of the fibres and release antimicrobial ions that penetrate the biofilm. Moreover, BG fibres can facilitate the direct release of ions into the biofilm matrix, ensuring a sustained antimicrobial effect. This dual-action capability not only disrupts the structural integrity of the biofilm but also ensures the continuous delivery of antimicrobial agents, enhancing the overall efficacy of the treatment. The insights gained from our mathematical model highlight the importance of these dual mechanisms and can be used to further optimise the design and doping of BG fibres to maximise their antimicrobial properties.

While our initial model provided valuable insights, future iterations will focus on incorporating spatial and temporal variability of EPS within the biofilm. With this we can better understand how EPS dynamics influence biofilm height and treatment efficacy. Specifically, exploring strategies for EPS removal could potentially enhance treatment outcomes by improving antimicrobial access to bacterial cells embedded within the biofilm matrix. BG fibres can be doped with antimicrobial ions as well as additional ions, such as copper, that target and remove EPS [49]. This multi-functional doping capability enhances the viability of BG fibres as a treatment option, as it allows for a thorough approach to disrupting the biofilm structure and improving the overall effectiveness of the treatment.

In addition, the current model considers only a single bacterial species, whereas clinical biofilms are frequently polymicrobial. Future work should therefore extend the framework to incorporate multiple interacting bacterial species with differing susceptibilities to antimicrobial treatment, enabling the investigation of how species composition and interspecies interactions influence biofilm development and treatment efficacy.

In conclusion, this study is the first to investigate the potential of BG fibres doped with antimicrobial ions as a treatment strategy against biofilm infections using mathematical modelling. This highlights the possibility of integrating BG fibres into clinical settings for biofilm-based wound care, which could help treat chronic wounds and reduce the burden on global health services. By integrating experimental data and refining our model parameters, we have gained valuable insights into the complex interplay of antimicrobial mechanisms within biofilms, predicting that treatment design should prioritise minimising the loss of ions over enhancing their diffusion into the biofilm. Moving forward, this simplified framework provides a foundation for incorporating additional biological complexity, including EPS dynamics, to better understand how these processes influence biofilm behaviour and treatment efficacy. Ultimately, when combined with experimental data, we can present pathways to optimise this therapeutic strategy aimed at combating biofilm-associated infections.

## Acknowledgments

Support from the Biotechnology and Biological Sciences Research Council (www.ukri.org/councils/bbsrc/) is greatly acknowledged, through the Midlands Integrative Biosciences Training Partnership (MIBTP) and research grant UKRI2819. We would also like to acknowledge support from the Engineering and Physical Sciences Research Council (grant code: EP/V051342/1, www.ukri.org/councils/epsrc/). SJ thanks the Leverhulme Trust (RPG-2019-382). We would like to thank Christopher Stark, University of Birmingham, for collecting the ICP-ms data. Finally we would like to thank Professor Paul Williams, University of Nottingham, for providing the bacterial strain, PAO1-N. The funders had no role in study design, data collection and analysis, decision to publish, or preparation of the manuscript.

